# An isoform-resolution transcriptomic atlas of colorectal cancer from long-read single-cell sequencing

**DOI:** 10.1101/2023.04.21.536771

**Authors:** Zhongxiao Li, Bin Zhang, Jia Jia Chan, Hossein Tabatabaeian, Qing Yun Tong, Xiao Hong Chew, Xiaonan Fan, Patrick Driguez, Charlene Chan, Faith Cheong, Shi Wang, Bei En Siew, Ian Jse-Wei Tan, Kai-Yin Lee, Bettina Lieske, Wai-Kit Cheong, Dennis Kappei, Ker-Kan Tan, Xin Gao, Yvonne Tay

**Affiliations:** Computer Science Program, Computer, Electrical and Mathematical Sciences and Engineering Division, King Abdullah University of Science and Technology (KAUST), Thuwal, Saudi Arabia; Computational Bioscience Research Center, King Abdullah University of Science and Technology; Cancer Science Institute of Singapore, National University of Singapore, Singapore; Core Labs, King Abdullah University of Science and Technology, Thuwal, Saudi Arabia; Department of Pathology, National University Health System, Singapore; Department of Surgery, Yong Loo Lin School of Medicine, National University of Singapore, Singapore; Division of Colorectal Surgery, University Surgical Cluster, National University Health System, Singapore; NUS Centre for Cancer Research, Yong Loo Lin School of Medicine, National University of Singapore, Singapore; Department of Biochemistry, Yong Loo Lin School of Medicine, National University of Singapore, Singapore

**Author notes:** Correspondence should be addressed to Yvonne Tay, Xin Gao and Bin Zhang, Email: Yvonne Tay,; Xin Gao,; Bin Zhang. These authors contributed equally to this work (Zhongxiao Li, Bin Zhang, Jia Jia Chan).

**Keywords:** colorectal cancer, long-read RNA-seq, scRNA-seq, neoantigen

## Abstract

Colorectal cancer (CRC) is the second leading cause of cancer death worldwide. In recent years, short-read single-cell RNA sequencing (scRNA-seq) has been instrumental in deciphering tumor cell heterogeneities. However, these studies only enable gene-level expression quantification but neglect alterations in transcript structures, which arise from alternative end processing or splicing, and are frequently observed in cancer. In this study, we integrated short- and long-read scRNA-seq of CRC patient samples to build the first isoform-resolution CRC transcriptomic atlas. We identified 394 dysregulated transcript structures in tumor epithelial cells, including 299 resulting from various combinations of multiple splicing events. Secondly, we characterized genes and isoforms associated with epithelial lineages and subpopulations that exhibit distinct prognoses. Finally, we built an algorithm that integrated novel peptides derived from predicted ORFs of recurrent tumor-specific transcripts with mass spectrometry data and identified a panel of recurring neoepitopes that may aid the development of neoantigen-based cancer vaccines.

## Introduction

The heterogeneity of cells and their composition is critical for cancer progression, patient survival and response to therapy ^1,2^. Single-cell RNA sequencing (scRNA-seq) enables RNA expression profiling for thousands of genes in hundreds and thousands of cells in parallel to facilitate the comprehensive characterization of these variations. scRNA-seq has been performed for colorectal cancer (CRC) to investigate tumor cell heterogeneities, transformation states, spatial organizations and immune cell infiltrations ^3–6^. Based on the gene expression profiles in each cell, scRNA-seq can distinguish malignant from infiltrated cells and group them into subpopulations with distinct molecular properties. Furthermore, the varying compositions of each cell subpopulation for both malignant and infiltrated cells have been utilized to classify tumors into subtypes to predict prognosis and response to therapy ^7,8^.

To date, most scRNA-seq cancer studies only quantify expression at the gene level. However, the majority of human genes can generate multiple transcript isoforms, particularly in tumor cells in which widespread alterations in transcript structure arising from 5’ and 3’ ends, and alternative splicing (AS) differences are frequently observed ^9–11^. Recently, we reported that pervasive splicing within the 3’-untranslated region (3’ UTR) is upregulated in cancer to promote oncogene expression and tumorigenesis ^12^. These results demonstrate that in addition to gene expression, changes in transcript structures also contribute to cancer pathogenesis. Traditionally, bulk RNA sequencing (RNA-seq) detects cancer-associated aberrations in specific regions of transcripts reconstructed from short reads, whereas the third-generation long-read sequencing (LR-seq), such as PacBio Iso-Seq, is able to directly capture dysregulations of full-length transcripts. Recently, two studies employed PacBio Iso-Seq with bulk RNA from breast cancer samples and gastric cancer cell lines to characterize their full-length transcripts, revealing extensive aberrant transcript structures that may be involved in tumor-related dysfunctions ^13,14^. However, only a handful of studies have combined LR-seq and scRNA-seq to dissect tumor cell heterogeneities and transcript dysregulations at the isoform level.

In addition, tumors specifically express RNA with dysregulated transcript structure (DTS) that may contain new open read frames (ORF) encoding unique proteins that serve as a potential source of neoantigen in tumor cells ^15^. Several studies have reported that tumor-associated intron retention, neo-splice junction, exitron and RNA editing could generate neoantigens ^9,16–18^. To date, only neoantigens derived from DNA mutations with the potential to activate the immune response have been used to generate personalized neoantigen vaccines in clinical trials for melanoma and glioma treatment ^19,20^. Studies for neoantigens arising from aberrant RNA processing are limited and primarily based on short-read RNA-seq data that rely heavily on annotations that could compromise the precision of full ORF predictions. Conversely, the identification of full-length tumor-specific anomalous transcripts using LR-seq may enable predictions of complete novel ORFs to more accurately and exhaustively derive neoepitopes. Nevertheless, this concept and its feasibility have not been explored.

Here, we present a study integrating short-read and PacBio Iso-Seq-based long-read scRNA-seq of matched CRC clinical samples to build a full-length transcriptomic atlas and investigate isoform-specific dysregulations in CRC. We identified 394 DTSs from 273 genes in the tumor epithelial cells, many of which were caused by the coupling of multiple splicing events. Additionally, we detected tumor cell-associated and isoform-specific RNA editing events. By classifying normal (EpiN) and tumor epithelial (EpiT) cells into subpopulations, we unveiled genes and transcripts associated with three EpiN lineages, as well as three EpiT subpopulations comprising more than 90% of malignant cells, each linked to a distinct prognosis. Finally, we predicted the proteome of CRC based on the full-length transcriptome. By integrating the novel ORFs from recurrent tumor-specific transcripts and mass spectrometry (MS) data, we developed an algorithm to build a panel of 22 neoepitopes with strong binding affinities to MHC molecules for a wide range of CRC patients, which may be exploited for the development of universal neoantigen-based cancer vaccines.

## Results

### Matched long- and short-read scRNA-seq atlas of human CRC

To study the landscape of full-length transcript isoforms in human CRC, we isolated single cells from eight tumor and eleven adjacent normal samples derived from 12 CRC patients, generated cDNA libraries using the 10x Genomics droplet-based method and performed matched Illumina short-read sequencing and PacBio Iso-Seq (**Fig. 1A, Extended Data Fig. 1a, Supplementary Table 1**). We curated a total of 18,966 high-quality cells with Illumina scRNA-seq data after applying conventional filtering (**Methods**, **Extended Data Fig. 1b**) with the number of cells per sample being slightly lower but comparable to related studies (**Extended Data Fig. 1c**) ^5,6^. To accurately identify the cell types, considering the lower cell number in our dataset, we did not perform clustering from scratch. Instead, we mapped these cells to the Human Colon Cancer Atlas (c295), which is currently the most comprehensive single-cell atlas of CRC ^6^. By integrating these two datasets (**Extended Data Fig. 1a**, **Methods**), the cells were grouped into 20 major clusters (**Fig. 1b**). We found that cells from the in-house tumor and normal samples were embedded into the c295 cell clusters without obvious batch effects (**Fig. 1c**). Sixteen out of 20 cell types from c295 were detected in the in-house dataset, of which eight consisted of more than 100 cells (**Supplementary Table 1c**). Among these, most of the cells were epithelial cells, followed by the immune and stromal cells (**Fig. 1d**), and the gene expression of representative cell markers were validated in each cell type (**Extended Data Fig. 1d-f**).

**Figure 1.**
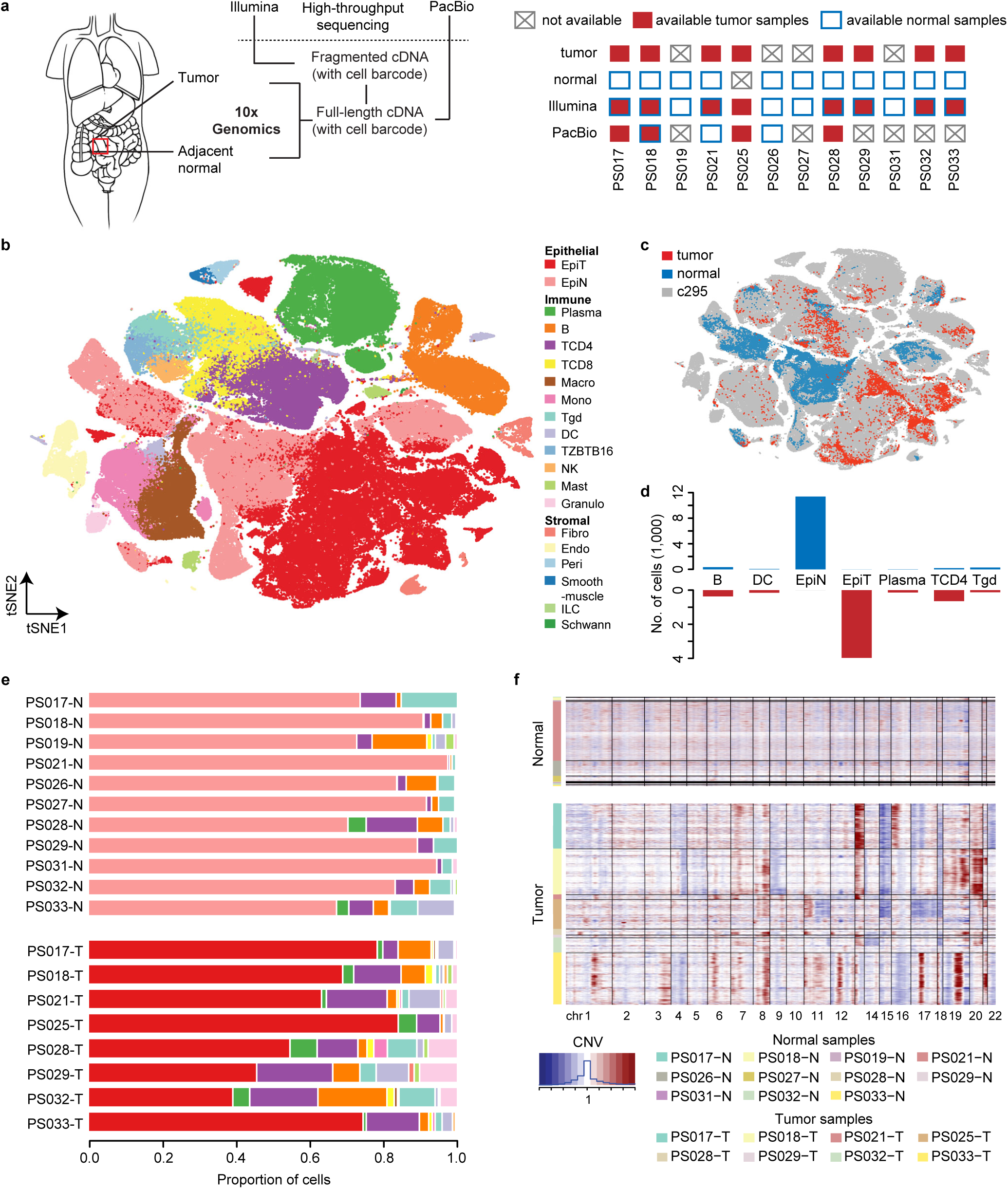
The short-read single-cell transcriptomic atlas of human CRC. (a) Schematic illustration of the workflow of long-read (PacBio) and short-read (Illumina) single-cell RNA sequencing (scRNA-seq) on tumor and normal samples from 12 CRC patients. (b and c) tSNE (t-distributed stochastic neighbor embedding) plots illustrating (b) the major cell types and (c) the source of cells from in-house tumor and normal samples, and the public dataset (c295). (d) Number of detected cells for each major cell type by in-house Illumina scRNA-seq data. (e) Proportion of detected cells from each major cell type in each sample. (f) Copy number variation (CNV) profiling of each cell from each sample.

Interestingly, compared to the normal samples, the tumor samples contained a higher number and proportion of immune cells, suggesting higher immunological activity in the tumor microenvironment (**Fig. 1e**, **Supplementary Table 1d**). To ensure robust identification of malignant epithelial cells, we inferred somatic copy number variations (SCNVs) of the cells from the normal and tumor samples using inferCNV ^21^. In general, the normal samples did not show significant SCNVs across the chromosomes, whereas the tumor samples displayed SCNVs in multiple regions (**Fig. 1f**). These regions were generally consistent within each sample and located within previously characterized cytobands, such as amplifications in 1q, 7p, 8q, 19q, 20p and 20q, as well as deletions in 4q, 15q and 22q ^22,23^. Using the inferred SCNVs and gene expression profiles (**Methods**), we obtained a collection of 3,943 and 11,388 reliable tumor (EpiT) and normal (EpiN) epithelial cells, respectively.

In addition to short-read scRNA-seq, we also performed long-read scRNA-seq on three normal and four tumor samples using PacBio Iso-Seq (**Fig. 1a**). Among the derived 19,264,405 circular consensus long-read sequencing (CCS) reads, 11,481,904 (59.6%) full-length non-concatemer (FLNC) reads were obtained (**Methods**). We identified 125,205 unique transcript isoforms from 17,753 genes, of which 23,379 were detected in both the normal and tumor samples. Based on the comparison of the splice junctions (SJs) of each aligned isoform with the reference transcriptome (GENCODE v37 and NCBI RefSeq release 109), we classified each transcript into four main categories: full splice match (FSM), incomplete splice match (ISM), novel in catalog (NIC) and novel not in catalog (NNC), as well as several other non-specific categories (antisense, genic, intergenic, fusion, etc.). FSMs and ISMs had fully or partially matched SJs against the reference, whereas the NIC and NNC transcripts contain novel SJs from a combination of known and novel splice sites, respectively (**Fig. 2a**). In total, we identified 31,837 (25.43%) FSM, 58,570 (46.78%) ISM, 13,291 (10.62%) NIC, 18,644 (14.89%) NNC and 2,863 (2.29%) isoforms in the other categories (**Extended Data Fig. 2a** and **2b**).

**Figure 2.**
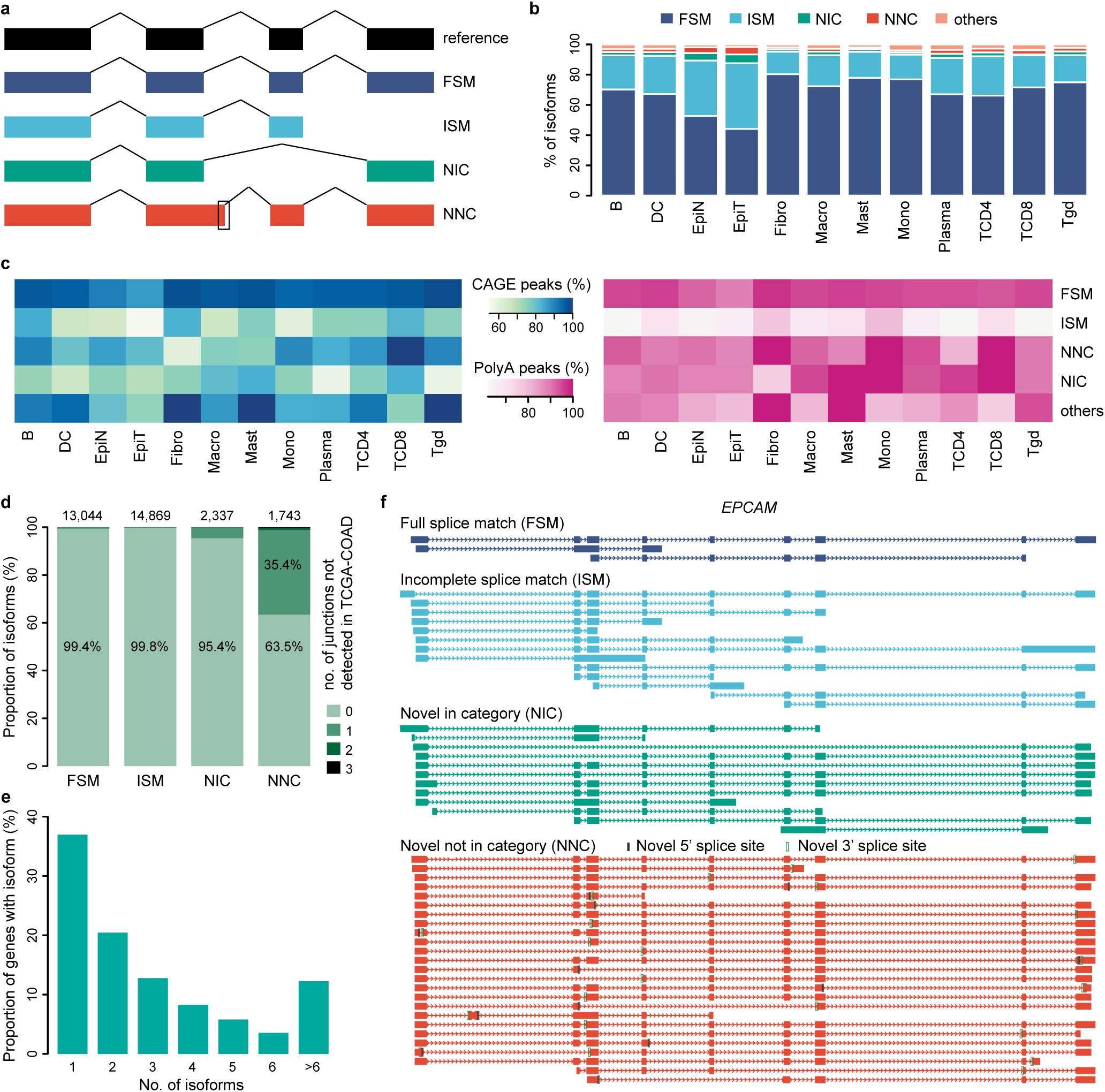
The long-read single-cell transcriptomic atlas of human CRC. (a) Schematic showing four structural types of isoforms identified by PacBio scRNA-seq data. FSM: full splice match, ISM: incomplete splice match, NIC: novel in catalog, NNC: novel not in catalog. The black box denotes a novel splice site. (b) Proportion of the identified transcripts from each isoform structural type in each major cell type. Others include antisense, genic (from intronic regions) and intergenic transcripts. (c) Percentage of the identified isoforms with supporting Cap Analysis of Gene Expression (CAGE) peaks for the 5’ ends and PolyA peaks for the 3’ ends. (d) Percentage of the identified isoforms with different numbers (0,1,2,3) of splice junctions that are not detected in the TCGA-COAD bulk RNA-seq data. (e) Percentage of genes with single and multiple detected isoforms. (f) Structure of the four types of isoforms from *EPCAM* identified by PacBio scRNA-seq data. The identified novel splice sites are highlighted with black boxes.

With long-read scRNA-seq, we were able to perform the quantification of RNA expression at the isoform level in each cell. Overall, we matched 1,994 out of 2,197 cell barcodes by comparing those identified in long-read and short-read scRNA-seq (**Extended Data Fig. 2c**). These cells were from 12 cell types, of which EpiN and EpiT are the most abundant, while immune and stromal cells constituted only a small fraction of the cell population (**Extended Data Fig. 3a**). Across all cell types, most of the isoforms were either FSM or ISM (**Fig. 2b**). Interestingly, we found that EpiT expressed a slightly higher proportion of novel isoforms (NIC and NNC), suggesting more transcriptomic abnormalities in tumor cells.

We further evaluated the quality of the detected isoforms by validating their structural components. The 5’-end of >60% and 3’-end of >70% of the isoforms overlapped with annotated transcription start sites (TSSs) and termination sites (TTSs) in each cell type (**Fig. 2c, Methods**). All the splice junctions from >99% of FSM and ISM isoforms were also detected in colon adenocarcinoma patients from the cancer genome atlas (TCGA-COAD) (**Fig. 2d**). ∼5% and >30% of NIC and NNC isoforms, respectively, contained at least one splice junction that has not been identified in the TCGA-COAD RNA-seq data, suggesting that some novel splice junctions could be more effectively captured by LR-seq. To evaluate the concordance between the long- and short-read scRNA-seq data, we compared and observed a high overlap of the expressed genes identified in both datasets (**Extended Data Fig. 3b**). Furthermore, we observed a significant correlation of gene expression quantified by these two methods in both EpiN and EpiT (*r* = 0.77, *p* < 2.2 × 10^−308^ and *r* = 0.67, *p* < 2.2 × 10^−308^, respectively) (**Extended Data Fig. 3c**).

Among the 11,138 genes detected by long-read scRNA-seq, we found that 63% expressed multiple isoforms, with 13% having more than 6 isoforms (**Fig. 2e**). For example, we detected 53 isoforms for the epithelial cell surface marker gene *EPCAM*. In addition to the 16 known isoforms (FSM and ISM), we identified 12 NIC and 25 NNC isoforms with the latter arising from 11 and 17 novel 5’ and 3’ splice sites (**Fig. 2f**). These results indicate a high transcriptomic complexity in CRC tumors, which has been overlooked in previous studies.

### Dysregulation of transcript structure and RNA editing in tumor epithelial cells

To systematically investigate the dysregulation of RNA isoforms in CRC, we defined two types of dysregulations by comparing gene and isoform expression between EpiN and EpiT (**Fig. 3a**). Dysregulated gene expression (DGE) refers to genes that have significantly different numbers of overall transcripts between the two cell types, while dysregulated transcript structure (DTS) is defined as a multi-isoform gene that expresses significantly different proportions of its isoforms between EpiN and EpiT (**Methods**). The fold changes of the genes showing significant DGE as measured by the two sequencing methods are strongly correlated (r = 0.77), with only three genes (<1%) showing discordance (**Fig. 3b**). This high concordance indicates the reliability of gene expression quantification using PacBio long-read sequencing data. We further experimentally validated the top three DGE events, demonstrating the exclusive expression of *MMP7* and *REG3A* in CRC tumor samples, while *REG1A* was significantly more highly expressed in the tumor samples compared to the adjacent normal (**Fig. 3c**). In total, we identified 898 DGE events and 273 genes with DTS from the PacBio data (**Fig. 3d**). We observed more DGEs that were upregulated (747) than downregulated (151), suggesting high transcriptional activity in tumor cells (**Fig. 3d**). Interestingly, we found that a higher frequency of genes with DTS had DGE, while single-isoform genes tend to be less dysregulated (**Fig. 3d**). Gene ontology (GO) enrichment analysis of the genes with DTSs showed enrichment of pathways related to RNA regulation, such as mRNA binding, translational initiation and RNA catabolic process (**Fig. 3e**). This suggests potential autoregulation of these genes, which is frequently observed for RNA-binding proteins, particularly splicing factors ^24–26^. Several immunological pathways were also implicated, indicating that DTS may contribute to dysregulated immunological activity in tumor cells. For example, the proliferating cell nuclear antigen gene, *PCNA*, presented both DGE and DTS. As a cell proliferation marker ^27^, its expression was upregulated in EpiT and detected in a higher proportion of cells (**Fig. 3f**). The EpiN and EpiT cells preferentially utilized distinct alternative first exons to differentially express the three *PCNA* isoforms. Overall, isoform proportion changes accounted only for a small fraction of transcripts with differential absolute expression levels between EpiT and EpiN for all four isoform categories, suggesting that transcriptional activity may have a more profound impact on isoform expression (**Extended Data Fig. 4a**).

**Figure 3.**
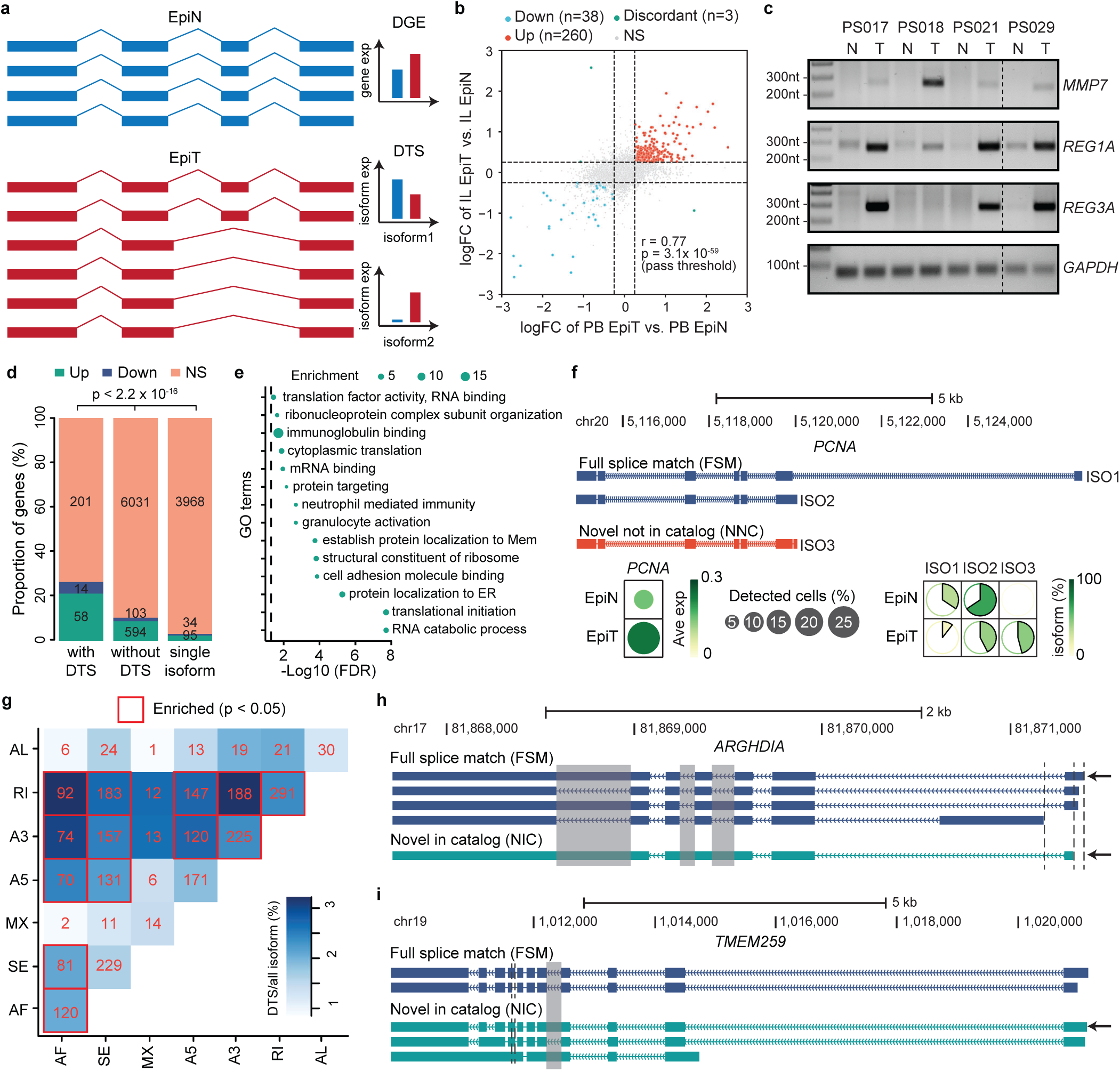
Dysregulated transcript structures in epithelial tumor cells. (a) Illustration of dysregulated gene expression (DGE) and dysregulated transcript structure (DTS) in epithelial tumor (EpiT) compared to normal (EpiN) cells. (b) Scatterplot showing the correlation in fold change of genes between EpiT and EpiN quantified by long- and short-read sequencing. Up (down): measured by both sequencing methods as significantly up- (down-) regulated; discordant: measured by both sequencing methods as significant but with inconsistent directions of change; NS: not significant. (c) PCR validation of the top three DGE events, *MMP7*, *REG1A* and *REG3A*, from (b) using four pairs of CRC tumor (T) and adjacent normal (N) patient samples (PS). (d) Proportion of upregulated (up), downregulated (down) and not significantly changed (NS) genes for those with DTS, without DTS and those with only one detected isoform. P-value from Chi-squared test. (e) Enrichment of Gene Ontology (GO) terms for genes with DTS. (f) Structures of the three identified isoforms from *PCNA* (upper), the gene expression pattern and the percentage of each isoform in EpiN and EpiT. (g) Numbers of DTS isoforms with co-occurrence of two types of AS events (or single type of events on the diagonal) and their percentage of the total corresponding isoforms. P-value from two-tailed binomial test. AF: alternative first exon, SE: skipped exon, MX: mutually exclusive exon, A5: alternative 5’ splice site, A3: alternative 3’ splice site, RI: retained intron, AL: alternative last exon. (h and i) Structural illustration of isoforms (arrows indicate DTS isoforms) from (h) *ARGHDIA* and (i) *TMEM259* with co-occurrence of two different types of splicing events. *ARGHDIA* DTS contains coupled AF and RI, while DTS from *TMEM259* contains coupled RI and A3.

We further applied SUPPA2 ^28^ to extract AS events for the identified transcript isoforms, which were categorized into alternative 3’ splice site (A3SS), alternative 5’ splice site (A5SS), alternative first exon (AF), alternative last exon (AL), retained intron (RI), skipped exon (SE) and mutually exclusive exon (MX) (**Extended Data Fig. 4b**, **Methods**). Consistent with previous observations ^13,14^, the most abundant AS events were from the AF, RI and SE categories (**Extended Data Fig. 4c**). We inspected how each isoform with DTS could arise from the different AS categories and found that isoforms derived from AF, A3 and RI were more enriched with DTS (**Fig. 3g**, diagonal). More importantly, our long-read scRNA-seq captured the structure of full-length isoforms, allowing us to investigate the coupling between different AS categories. We observed that the isoforms derived from coupled AF-RI, AF-A3 and A3-RI had the highest enrichment of DTS isoforms (**Fig. 3g**, off-diagonal). For example, among the *ARGHDIA* isoforms, those utilizing the distal first exon can retain any one of the three downstream introns (**Fig. 3h**), while the *TMEM259* isoforms using a proximal downstream 3’ splice site can retain an upstream intron (**Fig. 3i**). Our findings underline the unique advantage of LR-seq in capturing isoform complexity that have thus far eluded the short-read sequencing technology.

In addition to alternative splicing, post-transcriptional RNA modifications on single nucleotides could also impact the mRNA sequence and expression without altering transcript structures. Specifically, ADAR-mediated A-to-I RNA editing can modify both coding and non-coding regions of mRNAs, as well as long non-coding RNAs, and has been implicated in several autoimmune disorders ^29^ and multiple cancers ^30^.

We systematically identified the RNA editing events from isoforms sequenced using long-read scRNA-seq by counting the number of edited reads (**Methods**). To characterize the RNA editing level of a site, we computed RNA editing level per site (REPS) and per isoform (REPI), which are the ratio of the number of all edited reads against all reads for all isoforms and against all reads for a specific isoform, respectively (**Fig. 4a, Supplementary Table 2**). Of note, REPI calculation is only possible with LR-seq as it allows unambiguous association of each edited mRNA read with the isoform of origin.

**Figure 4.**
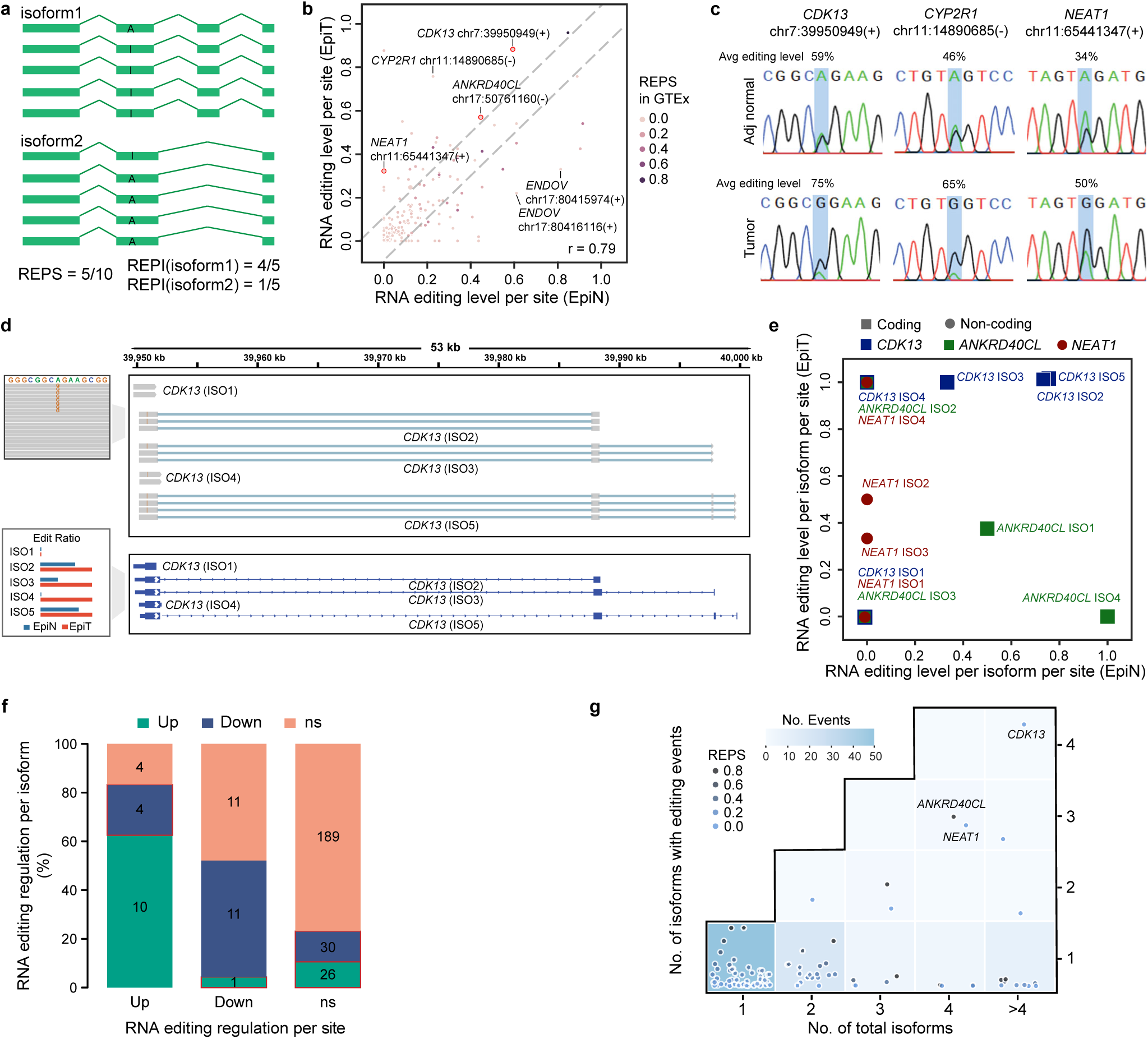
Dysregulated RNA editing in epithelial tumor cells. (a) Illustration of two approaches to calculate RNA editing levels: RNA editing level per site (REPS) and RNA editing level per isoform (REPI). (b) Scatter plot showing REPS for each detected event in epithelial tumor (EpiT) and normal (EpiN) cells. REPS based on RNA-seq data from GTEx are illustrated by scaled colors. (c) Sanger sequencing validation of the RNA editing sites and levels as shown in (b). Representative chromatograms are shown. (d) Illustration of the detected RNA editing events on each isoform from *CDK13* (upper) and the corresponding REPI in EpiN and EpiT. (e) REPI for each event on each isoform from the sites highlighted in (b) in *CDK13*, *ANKRD40CL* and *NEAT1*. (f) Consistency between REPS and REPI in EpiT compared to EpiN. Red boxes indicate inconsistencies between REPS and REPI. (g) Heatmap showing the number of isoforms with detected RNA editing versus the total number of detected isoforms of the gene. Each dot represents an editing event at a gene locus. Dots are colored according to their REPS and those for the three RNA editing events in Figure 4E are highlighted.

A total of 196 RNA editing sites with sufficient sequencing read support were identified in EpiN and EpiT cells. Most of these sites were on non-coding transcripts or the untranslated regions of coding transcripts, with more events in the 3’UTRs than 5’UTRs (**Extended Data Fig. 5a**), which is consistent with previous studies ^31–33^. Furthermore, 10 sites were found in the coding regions, which could potentially introduce codon alterations. For example, the tumor-associated event at chr7:39950949(+) that results in the amino acid change Q103R in CDK13 (**Fig. 4b**), was also observed in hepatocellular carcinoma ^34^. Based on the REPS, RNA editing events in genes such as *CDK13*, *CYP2R1* and *NEAT1* were significantly upregulated in EpiT compared to EpiN (**Fig. 4b**). We validated these events using normal colon, CCD 841 CoN, and CRC cell lines, DLD-1 and HCT116, as well as patient samples. The RNA editing levels for *CDK13, CYP2R1* and *NEAT1* were variable and did not follow any trend in the cell lines (**Extended Data Fig. 5b**). However, these were consistently higher in the patient tumor samples of at least four out of five matched pairs tested (**Fig. 4c** and **Extended Data Fig. 5c**), highlighting the importance of performing transcriptomic and validation studies using clinical samples.

To investigate isoform-specific RNA-editing, we computed and compared REPI between EpiN and EpiT. For example, we first extracted the PacBio long reads belonging to each *CDK13* isoform and calculated the editing level of the site on each isoform separately using the corresponding reads (**Fig. 4d**). We observed heterogeneity of RNA editing levels of the same site among different isoforms (**Fig. 4e** and **Extended Data Fig. 5d**). Four of the *CDK13* isoforms showed different editing levels in EpiN cells but consistently high editing levels in EpiT, whereas all four isoforms of *NEAT1* showed low editing levels in EpiN with varying editing levels in EpiT. Overall, the direction of RNA editing changes between EpiT and EpiN was largely concordant using REPS and REPI (**Fig. 4f**). However, due to lower read coverage, we only observed a maximum of four edited isoforms per gene for a specific editing site while the per-isoform RNA editing events were only observed on one isoform for most genes (**Fig. 4g**). Taken together, our results suggest isoform specificity of the RNA editing mechanism, which is still poorly understood.

### Genes and isoforms associated with normal epithelial cells from different lineages

As epithelial cells constituted the majority of the identified cells in the scRNA-seq profiles, we performed cell subtyping for them using the c295 dataset as a reference to investigate their subpopulations ^35^ (**Methods**). We identified 11 cell subtypes containing three differentiation lineages from the EpiN cells, including the newly characterized *BEST4+* lineage. The projection of the EpiN cells displayed a strong semantic structure in the two-dimensional tSNE space, suggesting diverse differentiation lineages of the subtypes (**Fig. 5a**). An increased level of differentiation was observed from the center to the edge in the tSNE space. For example, along the goblet lineage, cE02, cE06 and cE08 successively showed increased goblet maturity. Different combinations of subtypes were detected in each normal sample and the proportions were similar across multiple samples (**Fig. 5b**).

**Figure 5.**
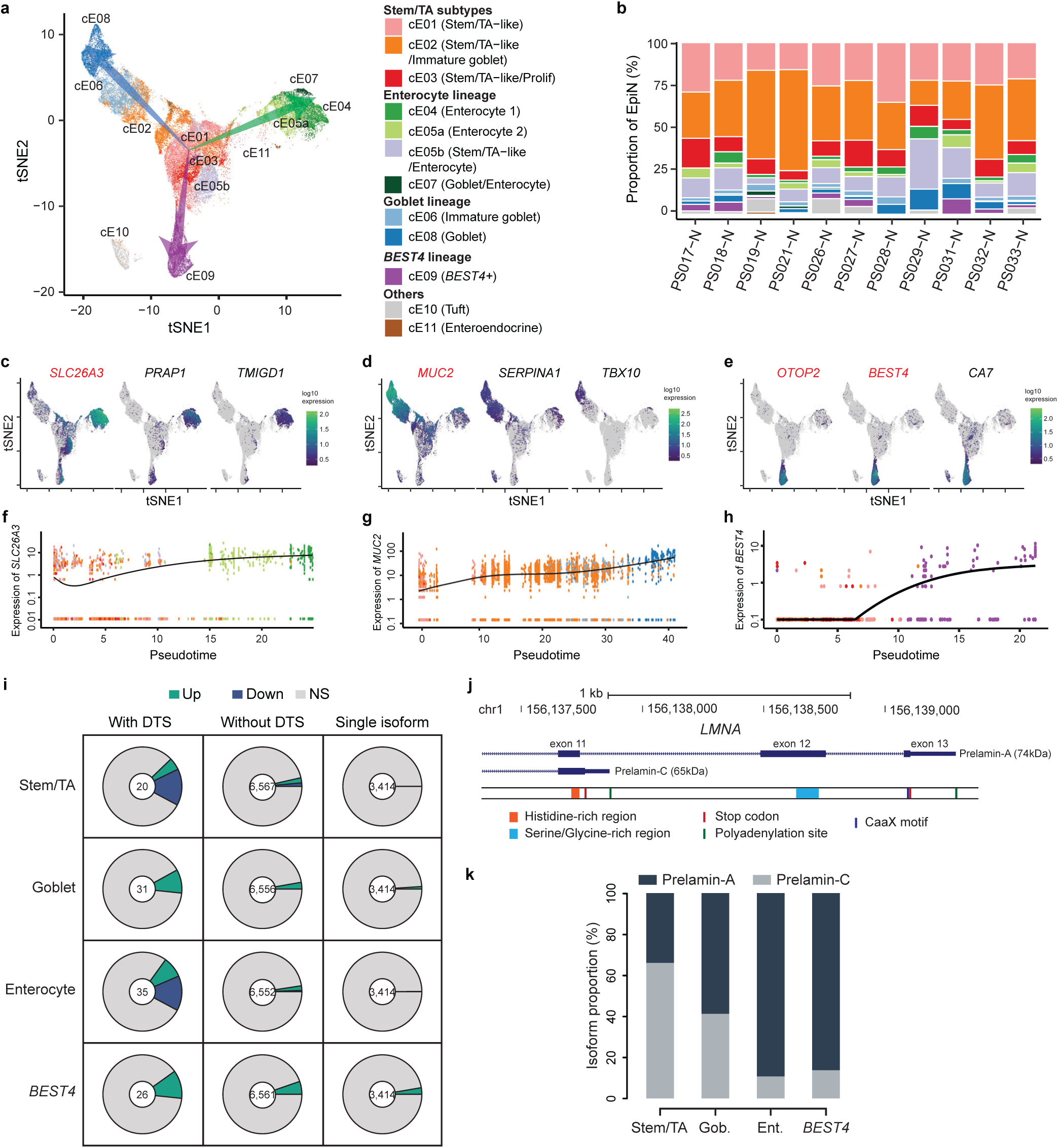
Transcriptome profiling of normal epithelial cell subtypes from multiple lineages. (a) tSNE (t-distributed stochastic neighbor embedding) plot illustrating subtypes of epithelial normal cells (EpiN) and the three main lineages of differentiation (indicated by arrows), including enterocytes (green), goblet (blue) and *BEST4* (purple). (b) Proportion of EpiN subtypes in each sample. (c-e) The top identified markers with lineage-specific expression for (c) enterocytes, (d) goblet and (e) *BEST4*. Known markers are in red. (f-h) RNA expression of lineage-specific marker genes over the pseudotime was estimated by trajectory analysis along the (f) enterocyte, (g) goblet and (h) *BEST4* lineages. (i) Proportion of upregulated (up), downregulated (down) and not significantly changed (NS) genes for genes with DTS, without DTS and those with only one detected isoform in the stem/TA subtypes and each differentiated lineage. (j) Structure of the *LMNA* transcripts encoding two Prelamin protein isoforms, Prelamin-A and Prelamin-C. (k) Proportion of the *LMNA* transcripts encoding the two Prelamin protein isoforms in the stem/TA subtypes and the three differentiation lineages.

Using monocle 3 ^36^, we inferred diffusion pseudotime along the differentiation trajectory of the three epithelial lineages (**Methods**) and identified 46, 31 and 38 lineage-specific genes (moran’s I > 0.5, adjusted p-value < 1.0 × 10^−5^) for the enterocyte, goblet and *BEST4* lineages, respectively, including both well-known and novel markers (**Fig. 5c-e**). These showed a steady increase in expression along the lineage with respect to pseudotime (**Fig. 5f-h**). We next performed differential gene and isoform expression analysis between the stem/transient amplifying (TA) and differentiated subtypes along the three lineages. Overall, we observed a similar number of up- and downregulated genes in the stem/TA subtypes, but mostly upregulation of lineage-specific genes in the three differentiated lineages (**Extended Data Fig. 6a** and **6b, Supplementary Table 3**). The genes with differential regulation were almost mutually exclusive between the three differentiated subtypes (**Extended Data Fig. 6c**), indicating distinct transcriptomic regulatory mechanisms to commit to a specific differentiation lineage.

By categorizing the genes in each differential gene and isoform analysis with DGE and DTS (defined in **Fig. 3a**), we observed a higher percentage of genes with DTSs subjected to DGE compared to those without DTSs or with only one isoform (**Fig. 5i**), similar to the trend observed between EpiN and EpiT (**Fig. 3b**). The gene *LMNA* encoding two protein isoforms, Prelamin-A and Prelamin-C, through AS and alternative polyadenylation (APA), serves as a representative of differential isoform usage between cell subtypes (**Fig. 5j**). We observed a trend of Prelamin-C to Prelamin-A switching when stem cells differentiated into enterocytes, goblet and *BEST4*+ cells (**Fig. 5k**), consistent with the findings in basal cell carcinomas, where Prelamin-A was negatively correlated to cell proliferation and appeared in the later stage of differentiation ^37^. Overall, our analysis provides the first comprehensive characterization of isoforms associated with three colon epithelial lineages.

### Tumor epithelial subtypes display different levels of stemness and differentiation

We subtyped the EpiT cells with the same classifier above to classify the EpiT cells to their closest EpiN subtypes (**Fig. 6a**). Consistent with previous reports ^6,7^, more than 93% of tumor epithelial cells were categorized as the three stem/TA subtypes (**Fig. 6b**), suggesting that malignant EpiT cells may possess stem/TA characteristics to maintain their proliferation and regeneration capabilities. Based on the integrative analysis of multiple CRC single-cell sequencing datasets, a recent study classified colon EpiT cells into two intrinsic consensus molecular subtypes (iCMS), iCMS2 and iCMS3 ^8^, and revealed that over 90% of EpiT cells are from either subtype. This holds true in ours and the c295 datasets with mostly one type of iCMS cells in each tumor (**Extended Data Fig. 7a** and **Extended Data Fig. 7b**). However, the distribution of the three subtypes from our categorization was more heterogenous in each tumor (**Fig. 6b** and **Extended Data Fig. 7b**). These distinct results suggest potentially diverse and independent molecular properties of the two classification methods.

**Figure 6.**
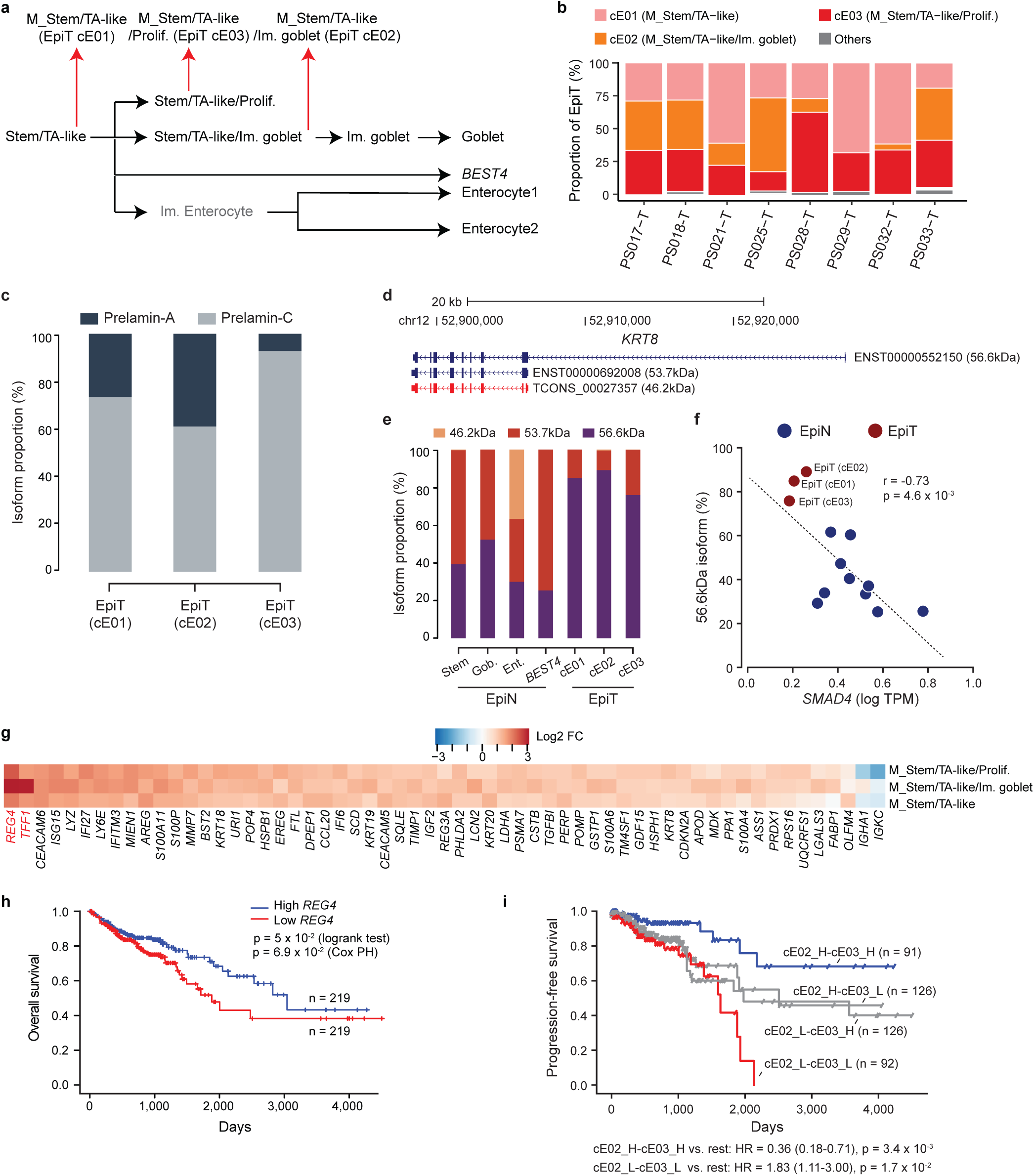
Dysregulation of genes and isoforms in the epithelial tumor cell subtypes. (a) Schematic showing the comparisons between each EpiT subtype and their corresponding EpiN cell subtype. (b) Proportion of EpiT subtypes in each sample. (c) Proportion of the *LMNA* transcripts encoding the two Prelamin protein isoforms in each EpiT subtype. (d) Structure of the *KRT8* transcripts encoding three CK8 protein isoforms with molecular weights of 56.6kDa, 53.7kDa and 46.2kDa. (e) Proportion of the *KRT8* transcripts encoding each CK8 protein isoform in each EpiN and EpiT subtype. (f) Correlation between the usage of the 56.6kDa transcript isoform and *SMAD4* expression in each EpiN and EpiT subtype. (g) Fold changes (log2 transformed) of top significant genes with dysregulated expression in each EpiT subtype compared to the corresponding EpiN subtype. (h) Overall survival of TCGA-COAD patients with different expression levels of *REG4*. (i) Progression-free survival of TCGA-COAD patients with different scores of cE02 and cE03 signature genes. H: high, L: low.

We further identified dysregulated genes in the EpiT subtypes by comparing them to the corresponding EpiN subtypes (**Extended Data Fig. 7c, Supplementary Table 4**). Most of these were shared among the three EpiT subtypes (**Extended Data Fig. 7d**), indicating a common underlying dysregulated mechanism of tumorigenesis. In addition to DGE, we performed differential isoform analysis for the EpiT subpopulations (**Supplementary Table 5**). Differential usage of the *LMNA* isoforms was also observed in the EpiT subtypes, with the highest usage of the Prelamin-C isoform in the most proliferative subtype, cE03, and Prelamin-A isoform in cE02 (**Fig. 6c**). Furthermore, we noticed that several cytokeratin genes, including *KRT8* and *KRT19*, were among the most significant DTS in all three EpiT subtypes (**Supplementary Table 5**). The overexpression of *KRT8* has been linked to increased tumor progression and invasiveness in several epithelial tumors ^38–40^. We detected three major *KRT8* isoforms, including two annotated and one novel isoform (**Fig. 6d**). The tumor subtypes mainly expressed the full-length isoform encoding a 56.6kDa protein, while the normal subtypes primarily utilized the short isoforms (**Fig. 6e**). Consistent with a previous study showing that the tumor suppressor *SMAD4* regulated *KRT8* splicing ^41^, we observed a negative correlation between *SMAD4* expression and the 56.6kDa isoform usage among the epithelial subtypes (**Fig. 6f**), further confirming the reliability of our isoform analysis. Overall, around 20 genes with DTS were identified in each EpiT subtype and consistent with those in the EpiN cells, genes with DTS were more likely to be DGE (**Extended Data Fig. 8a**). Contrary to the DGEs, the genes with DTS barely overlapped among the three subtypes (**Extended Data Fig. 8b**), suggesting potential dysregulation of isoform switching in the EpiT subpopulations.

Among the top dysregulated genes in the EpiT subtypes, *REG4* and *TFF1* showed the most striking upregulation in cE02 (**Fig. 6g**). Interestingly, we found that *REG4* and *TFF1* expression was highly correlated across the TCGA-COAD patients (r = 0.75, p = 3.81 × 10^−71^) and both were associated with better overall survival (OS) (**Fig. 6h** and **Extended Data Fig. 8c**). We speculate that the tumor samples with high *REG4* and *TFF1* expression may consist of a high proportion of cE02 cells, leading to better patient survival. To this end, we identified the marker genes in each subtype to define a signature score for the three subtypes (**Methods, Supplementary Table 5**). By classifying the TCGA tumors with cE02 and cE03 scores (**Extended Data Fig. 8d**), we found that the patient groups with both high cE02 and cE03 scores, and low cE02 and cE03 scores, had the best and worst OS and progression-free survival (PFS), respectively (**Fig. 6i** and **Extended Data Fig. 8e**), suggesting a synergistic effect of cE02 and cE03 scores on patient prognosis. Similar trends were observed using an independent microarray cohort curated by the CRC Subtyping Consortium (CRCSC) ^42^ (**Extended Data Fig. 8f** and **Extended Data Fig. 8g**). As cE02 and cE03 have relatively lower stemness features and higher proliferative activity compared to cE01, these results indicate that CRC tumors enriched with high stemness and low replicative cells are associated with more frequent progression and shorter OS time.

### Identification of neoepitopes from recurrent tumor-specific isoforms for cancer vaccine development

To investigate the effect of transcript structure alterations on the proteome, we predicted the open reading frames (ORFs) of all the identified isoforms. We performed a series of analyses on the predicted proteome and identified 8,984 novel ORFs from NIC and NNC isoforms (**Extended Data Fig. 9a-e**, **Supplemental Notes Section S1**). After integrating publicly available mass spectrometry (MS) data ^43^ from TCGA-COAD patients and in-house MS data from matched tumor and normal CRC patient samples and cell line, HCT116, 26 and 44 novel proteins from the NIC and NNC isoforms were supported (**Extended Data Fig. 9f**). There were also more MS-supported novel isoforms from EpiT and tumor samples compared to EpiN and normal samples, suggesting that transcriptomic dysregulation in tumor cells could result in increased abnormalities of the proteome (**Extended Data Fig. 9g**).

As abnormal proteins produced in cancer cells are critical for spontaneous anti-tumor immune response, they serve as a potential source for neoantigen-based cancer vaccine development to leverage the immune system for cancer treatment ^15^. Therefore, we systematically investigated putative neoepitopes in CRC derived from tumor-specific transcripts. Using the identified novel isoforms together with annotated isoforms as reference, we quantified the expression of each isoform in the TCGA-COAD samples (**Fig. 7a**, **Methods**). We retained 12 recurrent (>5 samples) novel isoforms with significantly higher amount of expression and number of supporting reads of novel junctions in the TCGA-COAD tumor samples than normal samples and whose ORF-derived peptides were MS-supported (**Fig. 7b**, **Methods**). Out of five isoforms selected for experimental confirmation, we validated the unique splice junctions and sequences for three isoforms arising from AF and A5SS AS events, and one with combined A5SS and retained intron (RI) in colon epithelial and CRC cell lines, as well as at least four of the five matched patient samples tested (**Fig. 7c, Extended Data Fig. 10a** and **10b, Supplementary Table 6**). All novel 9-mers derived from the predicted ORFs of the four experimentally validated isoforms were regarded as candidate neoepitopes in CRC.

**Figure 7.**
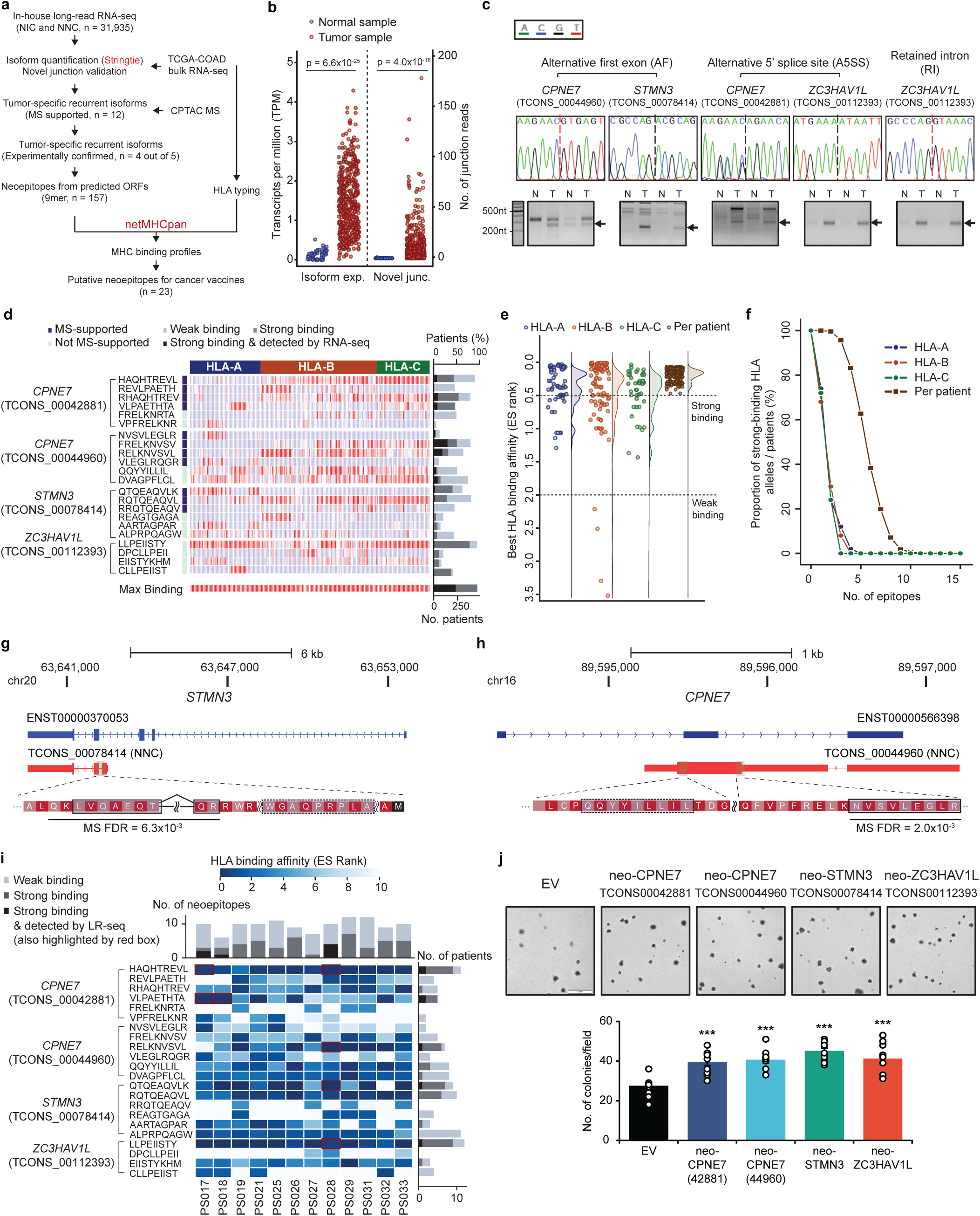
Identification of common neoantigens from recurrent tumor-specific transcripts. (a) Workflow for the identification of neoepitopes from novel tumor-specific recurrent transcript isoforms for cancer vaccine development. (b) Estimated maximum expression level (quantified by Stringtie) of MS-supported candidate isoforms and the number of supporting reads of novel junctions in the normal and tumor TCGA-COAD samples. (c) PCR and Sanger sequencing validation of the unique splice junctions for the recurrent tumor-specific isoforms from which the neoepitope panel is going to be derived. Novel splice junctions are depicted by black dotted lines and 5’ junction of sequences unique to selected isoforms by red dotted lines. Representative chromatograms and images are shown. (d) HLA binding profile of the panel of 22 neoepitopes against the HLA alleles (HLA-A, HLA-B, HLA-C) among the TCGA-COAD patients. The right panel summarizes the number of patients with at least one HLA allele showing ‘weak/strong binding’ affinities to the neoepitope of the corresponding row. ‘Strong binding & detected by RNA-seq’ denotes neoepitopes derived from isoforms with TPM>0.5 (as in Fig. 7B). The last row summarizes the percentage of TCGA-COAD patients with at least one neoepitope satisfying the two criteria above. (e) HLA binding profile of the selected neoepitopes. Each dot represents the best binding affinity of MHC molecule(s) from an HLA allele (columns 1-3) or a TCGA-COAD patient (column 4) against the panel of neoepitopes. Neoepitopes whose sequences overlap with MS are marked as ‘MS supported’ and vice versa. (f) Percentage of HLA alleles or patients with at least n (indicated by the x-axis) strong-binding neoepitopes in the panel. (g-h) Selection of neoepitopes for a novel isoform of (g) *STMN3* (TCONS_0078414) and (h) *CPNE7* (TCONS_00044960). The selected MS-supported neoepitopes are encircled by solid boxes and the non-MS-supported ones by dashed line boxes. (i) The binding affinity of the 22 neoepitopes for the 12 in-house patients. ‘Weak/strong binding’ denotes the binding affinity of the neoepitopes to at least one patient HLA allele (each row). ‘Strong binding & detected by LR-seq’ neoepitopes are indicated by red boxes in the corresponding heatmap. The top panel summarizes the total number of neoepitopes that satisfies each criterion above per patient. The right panel summarizes the total number of patients for whom the epitope satisfies the above criteria. (j) Effect of overexpressing the open reading frames derived from the validated neoepitopes in (c) on anchorage-independent growth in DLD-1 cells.

As a limited number of epitopes can be loaded into a cancer vaccine, it is imperative to maximize vaccine efficacy by selecting an optimized subset of neoepitopes with a high binding affinity to the class I MHC molecules that facilitate antigen presentation. To develop cancer vaccines for a broad range of patients, we considered the diversity of HLA alleles across different patients and predicted the MHC binding profiles of 458 TCGA-COAD patients to the neoepitopes from the four recurrent tumor-specific isoforms. We formulated a greedy algorithm to select four to six neoepitopes from each of the four isoforms to maximize their coverage to the HLA alleles or patients, while prioritizing those that overlapped with MS-supported peptides (**Fig. 7d**, **Extended Data Fig. 11a, Supplementary Table 7, Methods, Algorithm 1**).

Among the 22 selected neoepitopes, there were always one or more showing strong binding affinities (ES rank < 0.5) to 74% of HLA-A, 67% HLA-B and 70% HLA-C MHC molecules, with only four HLA-B alleles not passing the weak binding threshold (ES rank > 2.0) (**Fig. 7e**). For each patient, the highest binding affinities with their three types of MHC molecules were always above the strong binding threshold (**Fig. 7e**). All patients carry at least three strong-binding neoepitopes and 50% of them have at least five strong-binding neoepitopes from the panel (**Fig. 7f**). To better represent the global population, we generated another slightly different panel of neoepitopes (**Extended Data Fig. 11b, Supplementary Table 7**) from the same four isoforms but optimized for the top 50 high-frequency alleles in each HLA type ^44^, which similarly showed strong HLA binding affinities (**Extended Data Fig. 11c**).

To illustrate an example from our panel, the neoepitopes selected for the tumor-specific novel isoform of *STMN3* (TCONS_0078414) that utilizes a coding alternative first exon, were ‘RQTQEAQVL’ that spans the novel splice junction and overlaps with MS-detected peptides, and ‘ALPRPQAGW’ that has a strong HLA binding affinity (**Fig. 7d**). The neoepitopes for *CPNE7* resulting from an alternative 5’ splice site were similarly selected (**Fig. 7h**). The HLA binding profile of the neoepitope panel showed heterogeneous affinities to the three types of HLA (**Fig. 7d**). Overall, all patients have at least one strong-binding neoepitope and 50.4% (n=231) of them with corresponding isoform detection in the tumor samples (**Fig. 7d**). Furthermore, using the same panel of neoepitopes optimized for TCGA patients, we also observed strong HLA binding potential in the 12 in-house patients, with all patients having at least six strong-binding neoepitopes (**Fig. 7i, Supplementary Table 8**). In addition to serving as a source of neoantigen, we speculate that these tumor-specific isoforms may also have certain oncogenic functions considering their tumor-specific expression pattern. In line with this, we found that the overexpression of their ORFs promotes cell growth *in vitro* (**Fig. 7j** and **Extended Data Fig. 10c, d**). Collectively, these findings demonstrate a proof-of-concept for the discovery of neoepitopes based on the tumor-specific transcriptome, as well as the invaluable application potential of long-read scRNA-seq in the development of neoepitope-based cancer vaccines.

## Discussion

The intestinal epithelium is the fastest self-renewing tissue in mammals, which contains epithelial cells with diverse differentiation statuses, proliferation activities, and other properties and functions. Despite various markers and gene expression profiles that have been characterized for each epithelial cell subtype, the diversity of RNA isoforms in these subtype remains underappreciated. Our study provides a full-length transcriptome across different epithelial subtypes and identifies isoforms associated with enterocyte, goblet and *BEST4* lineages, respectively. Although alternative RNA isoforms have been shown to regulate cell fate determination of stem cells ^45^, the prevalence and potential role of alternative isoforms in intestinal epithelial differentiation are poorly understood, therefore our data may provide a valuable resource to facilitate further investigations.

CRC is the third most common cancer type and ranks as the second leading cause of cancer death around the world ^46^. Consistent with the first high-throughput scRNA-seq study on CRC and other single-cell analyses that traced the transformation of polyps to CRC ^3,4^, we find that the vast majority of CRC tumor cells have stem-like cell characteristics. However, studies for EpiT subtypes are limited. Based on the integrative analysis of multiple CRC single-cell sequencing datasets, a recent study classified colon EpiT cells into two subtypes and showed that 90% of EpiT in each tumor were from either subtypes ^8^. They further found that these two subtypes were associated with different cancer driver mutations, suggesting their classification method mainly captured heterogeneities across different tumors. However, our EpiT subtype classification demonstrated that each tumor comprised a substantial proportion of three cell subtypes with varying differentiation status and proliferation activities, thus capturing more intratumor heterogeneities. As our classification indicates that CRC tumors are differentially associated with PFS and OS depending on their subtype composition, the markers we propose for each subtype may serve as useful prognostic indicators. In addition to epithelial cells, other stromal and immune cells may also impact patient survival and response to treatment ^5,7^. However, due to limited cell numbers, our study could not effectively capture the full-length transcriptome in these cells, highlighting the need for more advanced and robust cell isolation technologies.

Despite recent reports of the full transcriptome by bulk RNA LR-seq in breast and gastric cancers ^13,14^, single-cell LR-seq studies are rare and limited to cancer cell lines ^47,48^. Moreover, previous LR-seq studies in cancer focused only on alterations in single AS events. Here, we identify hundreds of DTS arising from combined AS events and calculate isoform- and cell-type-specific RNA editing levels, all of which originate from primary CRC tissues and are only achievable by single-cell LR-seq. Nonetheless, our analyses are limited by technological constraints leading to the relatively low coverage of single-cell PacBio sequencing. Thus, the development of methods to increase the capture efficiency of single-cell isoform sequencing will be beneficial for future transcriptomic studies ^49^.

Therapeutic neoantigen-based cancer vaccines aim to leverage the immune system for the treatment of cancer ^15^. Due to somatic mutations and transcriptomic dysregulation, tumor cells produce abnormal proteins that are absent in normal cells. In the event that fragments of these proteins are transported to the cell surface via the MHC I pathway, they may potentially be recognized by the immune system as an antigen. We show that the full-length transcriptome in tumor cells with MS data from matched patient samples can provide a comprehensive source to detect aberrant mRNA-derived neoantigens. Moreover, we propose an algorithm that provides an optimized combination of recurrent tumor-specific neoepitopes with high HLA binding affinities for a wide spectrum of CRC patients. Recently, the phase II clinical trial for a neoantigen-based mRNA vaccine by Moderna has met the endpoint ^50^. In contrast to the customized vaccine designed based on a neoepitope identified by exome sequencing of a patient tumor, the panel we propose can potentially target larger CRC patient cohorts without sequencing individual tumors. This could facilitate the development of neoantigen-based cancer vaccines with less economic and time constraints to benefit a larger population.

## Methods

### CRC clinical samples

The protocols for the human studies comply with all relevant ethical regulations and are approved by the National Healthcare Group Domain Specific Review Boards (NHG DSRB Ref: 2018/01032). All patients gave informed written consent. The CRC tissue samples were derived from 12 patients from National University Hospital (details on the patients and samples are summarized in **Supplementary Table 1**).

### Tissue processing, library construction and single-cell sequencing

Colon tissues from patients were cut into approximately 5 mm pieces and washed three times with cold PBS until the supernatant became clear. The supernatant was discarded and the tissue pieces were further cut into 1-2 mm pieces. 5 ml of PBS containing 5 mM EDTA (EDTA-PBS) was added to the tissue fragments and incubated at 4°C for 30 minutes with gentle agitation, followed by centrifugation at 200 *x*g, 4°C for 5 minutes. The supernatant was discarded and the tissue fragments were vigorously resuspended in 5 ml of cold EDTA-PBS. The fragments were allowed to settle by gravity and the supernatant was collected in a fresh 15 ml tube. This resuspension/ sedimentation step was repeated five times. The supernatant from each round was collected in a separate 15 ml tube and checked under an inverted microscope for single cells. All fractions containing single cells were combined and the cells were pelleted by centrifugation at 200 *x*g, 4°C for 5 minutes, and washed with cold EDTA-PBS. The single cells were resuspended in serum-free DMEM containing 0.05% trypsin and 5 U/ml DNaseI and incubated at 37°C with shaking for 15 minutes. The dissociated cells were centrifuged at 200 *x*g, 4°C for 5 minutes, the supernatant was removed and the pellet was resuspended in 2 ml of serum-free DMEM supplemented with 2 U/ml DNaseI, and filtered using a 35 mm mesh. The cells were pelleted and washed once with cold PBS and resuspended in PBS supplemented with 0.4% BSA, followed by cell quantification.

For GEM generation, 10,000 cells from each tissue sample were used to load the Chromium Next GEM Chip G (10x Genomics) and Chromium controller following the manufacturer’s protocol (Chromium Next GEM Single Cell 3’ v3.1). Post GEM-RT cleanup, cDNA amplification and library construction were performed according to the manufacturer’s instructions with some modifications. Prior to the fragmentation step, 20 µl of cDNA from each sample were separately aliquoted for PacBio long-read sequencing. To generate the Single Cell 3’ Gene Expression library for Illumina sequencing, 10 µl of cDNA were processed as per the 10x Genomics protocol, with an extension time of 3 minutes in the sample index PCR step. The final Illumina libraries were sequenced on the NovaSeq 6000 platform with S1 or S4 flowcells using the recommended sequencing protocol.

The remaining full-length cDNA from the 10x Single Cell workflow was processed for PacBio long-read sequencing. The standard PacBio protocol for single cell Iso-Seq libraries was used with the minimum number of PCR cycles (typically 12 cycles) for amplification of cDNA to increase the mass of the input DNA prior to starting library preparation. Following purification, the library was processed with the normal DNA damage and end-repair steps, and SMRT Bell adapter ligation. The final library was then bound, loaded onto 8M SMRT cells and sequenced on the PacBio Sequel II instrument with a 24-hour movie time.

### Plasmids

The open reading frames (ORFs) of validated neoepitopes followed by a HA tag were cloned into pcDNA3.1+ using the primers and restriction sites listed in **Supplementary Table 9**. All constructs were verified by Sanger sequencing.

### Cell culture and transfection

Human normal colon cell line, CCD 841 CoN (ATCC: CRL-1790), was cultured in Dulbecco’s Modified Eagle Medium (DMEM). Human CRC cell lines, DLD-1 (Horizon Discovery: HD PAR-086) and HCT116 (ATCC: CCL-247) were cultured in Roswell Park Memorial Institute (RPMI) 1640 Medium and DMEM, respectively. All culture media were supplemented with 10% FBS, glutamine and penicillin/streptomycin. The cells were maintained at 37°C and 5% CO_2_ in a humidified atmosphere. For overexpression experiments, cells were seeded at 100,000 cells per well in 12-well plates 24 h prior to transfecting 1 µg of each plasmid using Lipofectamine 3000 (Thermo Fisher) following the manufacturer’s instructions.

### Soft agar assay

Cells were transfected 18-24 h prior to seeding as described above. A 0.6% base agarose was prepared in 12-well plates on the day of seeding. Transfected cells were trypsinized, harvested and counted. Seeding densities of 5,000 and 7,000 cells per well were used for CCD 841 CoN and DLD-1, respectively. The cells were resuspended in their respective growth media, mixed with agarose to a final agarose concentration of 0.3% and seeded on the prepared base. After agarose solidification, 0.5 ml of growth medium was added to each well. The cells were maintained in conditions described above and the growth medium was changed every 2-3 days. The colonies were imaged after 7-14 days under 4x magnification using the Olympus IX71 microscope and quantified using ImageJ (v1 51j8).

### RNA extraction and PCR

Trizol and the PureLink RNA Mini Kit (Thermo Fisher) were used to extract total RNA from DLD-1 and HCT116 cell lines. The ISOLATE II RNA Mini Kit (Bioline) was used to extract RNA from CRC patient samples following the manufacturer’s protocol. cDNA was generated using the High Capacity cDNA Reverse Transcription Kit (Thermo Fisher). PCR experiments were performed using Platinum™ Taq DNA Polymerase (Thermo Fisher). Subsequently, PCR products were subjected to agarose gel electrophoresis, gel extraction and purification using the QIAquick Gel Extraction Kit (Qiagen) and Sanger sequencing. Chromatograms were visualized and multiple sequence alignments were performed using SnapGene (v6.2.1). RNA editing levels were quantified using ImageJ (v1.50). PCR primers used are listed in **Supplementary Table 9**.

### Protein extraction and western blot analysis

Cells were harvested 48 h post-transfection, lysed and 10 µg of lysates were fractionated using 12% SDS-PAGE followed by transfer to PVDF membranes as described previously ^12^. Specific primary and secondary antibodies in 5% BSA-TBST were incubated with the membranes to probe for genes of interest.

### Mass Spectrometry

To generate an in-house mass spectrometry dataset of matched tumor-normal sample pairs, <10mg tissue for each tissue were processed with the iST sample preparation kit (Preomics). Samples were taken up in 100 μL LYSE buffer and mixed with 50 mg of protein extraction beads (Diagenode). Samples were sonicated with 10-20 cycles at 30sec ON/OFF in a Diagenode Bioruptor Plus until complete tissue disruption was observed. Protein concentrations were determined using the Pierce BCA Protein Assay Kit (Thermo Scientific). 20 μg of protein per sample were digested with the iST kit according to the manufacturer’s instructions and samples were eluted with the fractionation add-on into 3 fractions. Digested peptides were quantified with the Quantitative Fluorometric Peptide Assay kit (Thermo Scientific), normalized to 0.1 μg/μL and stored at -20°C prior to analysis on an EASY-nLC 1200 Liquid Chromatograph (Thermo Scientific) coupled to a timsTOF fleX (Bruker) mass spectrometer. For each sample, 5 μL digested peptides (500 ng) for each fraction were injected and separated on an Aurora series column (25 cm length, 75 μm inner diameter, C-18 1.7 μm; IonOpticks) with an integrated captive spray emitter. The column was mounted on a captive spray ionization source and temperature controlled by a column oven (Sonation) at 50°C. A 105-min gradient from 2 to 40 % (v/v) acetonitrile in 0.1 % (v/v) formic acid at a flow of 400 nL/min was used. Spray voltage was set to 1.65 kV. The timsTOF fleX was operated with data-dependent acquisition (DDA) in PASEF mode with 10 PASEF ramps per topN acquisition cycle (cycle time 1.17s) and a target intensity of 10,000. Singly charged precursor ions were excluded based on their position in the m/z-ion mobility plane and precursor ions that reached the target intensity were dynamically excluded for 24 seconds.

### Summary of bioinformatic analysis

The bioinformatic analysis workflow is summarized in **Extended Data Fig. 13**. Additionally, the command lines and parameters used in the analysis are also provided (**Supplemental Notes Section S2**).

### Illumina short-read scRNA-seq data pre-processing

CellRanger v4.0.0 (https://support.10xgenomics.com/single-cell-gene-expression/software/downloads/latest) was used for the pre-processing of 10x droplet-based scRNA-seq raw data. Specifically, ‘cellranger count’ with default parameters was used to align sequencing reads to the human reference genome (GRCh38). The gene expression count matrix was then obtained based on the genomic coordinates defined by the GENCODE annotation (release v37 for GRCh38). Subsequently, ‘cellranger aggr’ was performed to aggregate the count matrices of samples in multiple ‘cellranger count’ runs. The aggregated count matrix of all samples was then loaded into Seurat (v3.2.3) ^51^ for subsequent analysis. A cell was filtered out if it satisfied any of the following criteria: (1) expression of fewer than 250 genes, (2) detected fewer than 300 UMIs, (3) more than 70% of the UMIs mapped to mitochondrial genes. All mitochondrial genes were discarded in subsequent analysis. This analysis resulted in a total of 18,966 high-quality cells from 22 tissue samples.

The gene expression count matrix was log-normalized using Seurat’s ‘NormalizeData’ function. The top 2000 genes with the highest variations were selected using the ‘SelectIntegrationFeatures’ subroutine of Seurat, and the ‘IntegrateData’ subroutine was used to remove batch effects across the samples. Integration anchors were chosen based on the first thirty dimensions of the canonical correlation analysis (CCA) ^51^. We performed the analysis with above procedures separately on the normal and tumor samples to prevent the batch effect correction procedure from eliminating differences between the normal and tumor cells (**Extended Data Fig. 1**).

### Cell type identification

We integrated the in-house data to a reference dataset, Human Colon Cancer Atlas (c295) ^6^from the Broad Institute Single Cell Portal (https://singlecell.broadinstitute.org/), for more robust and reliable identification of the cell types. The c295 atlas contains gene expression profiles of 371,223 cells. Due to memory constraint, a subset of the atlas containing 50% of the cells in the c295 atlas was used as the reference based on a stratified sampling procedure across different cell types. Transfer anchors were chosen by the ‘FindTransferAnchors’ subroutine of Seurat using the first thirty dimensions of the principal component analysis (PCA). The ‘TransferData’ subroutine was used to transfer the c295’s ‘ClusterMidway’ annotation to the in-house data via the selected anchors. The ‘ClusterMidway’ labels of the c295 atlas annotate cells into 20 different types, including (with boldface indicating the availability of the cell type in the in-house data):

- 2 epithelial cell types: **normal epithelial cells (EpiN**) and **tumor epithelial cells (EpiT**).
- 3 immune cell types: **B cells, plasma cells, CD4+ T cells, CD8+ T cells, gamma-delta T cells, PLZF+ T cells, natural killer cells, innate lymphoid cells, dendritic cells, granulocytes, macrophages, mast cells, monocytes,**
- 5 stromal cell types: **fibroblasts**, endothelial cells, pericytes, smooth muscle cells, Schwann cells

Several post-processing steps were performed for the result from the above classification: (1) We corrected all the EpiN misclassified as EpiT cells in the normal samples. (2) To improve the purity of the identified epithelial tumor cells, we trained a gradient boosting classifier (xgboost v1.5.0 ^35^) using the combined EpiN and EpiT cells from the c295 atlas and the in-house dataset. Following a similar 5-fold cross-validation scheme as in ^6^, we divided the combined dataset into five independent splits and used four folds for training and the rest one fold for testing. The classifier was trained to perform the binary classification of whether a cell is a EpiT or a EpiN cell. Only when a cell came from a tumor sample and with >75% prediction probability as EpiT cells in the test split did we regard it as a candidate for the third step. (3) For all candidates from step (2), we inferred somatic copy number variations (SCNV) of the cells based on their gene expression profiles using inferCNV (v1.11.1, https://github.com/broadinstitute/infercnv) ^21^. The final set of tumor epithelial cells was kept as those having >10% genomic SCNV regions. The statistics of epithelial cells that passed the xgboost and the SCNV criteria are reported in **Supplementary Table 1e**. The epithelial cells from tumor samples that did not pass either the gradient boosting criteria or the SCNV criteria were deemed as undetermined epithelial cells.

### Dimensionality reduction and visualization

PCA was performed on the log-normalized and centered gene expression matrix for dimensionality reduction. t-distributed stochastic neighbor embedding (t-SNE) ^52^ was then performed to generate 2D embeddings for visualization. tSNE was supplied with the first 30 components of PCA and run with a perplexity value of 30.

### PacBio long-read scRNA-seq data pre-processing

SMRT Link (v9.0.0, https://www.pacb.com/support/software-downloads/) was utilized to pre-process PacBio Iso-seq long-read scRNA-seq data. We used the Isoseq-deduplication pipeline ^53^ recommended by the official repository of PacBio (https://github.com/PacificBiosciences/IsoSeq/) for the processing of raw data from the PacBio Sequel II system. The ‘ccs’ module of SMRT Link was applied to generate consensus reads (CCS Reads) from the BAM file that contained the PacBio subreads. The reads with less than 90% accuracy (‘--min-rq 0.9’) were discarded. The ‘lima’ module was utilized in orientation determination and primer removal of the reads. The cell barcodes and UMIs of each read were extracted by ‘isoseq3 tag’ using the library design ‘T-12U-16B’. The processed reads were trimmed of the Poly(A) tail and potential concatemers were removed by the command ‘isoseq3 refine’. The resulting full-length non-concatemer (FLNC) reads were clustered together according to their cell barcode and UMI with ‘isoseq3 dedup’ to deduplicate and generate a consensus sequence of each molecule. Those consensus sequences were then mapped to the human reference genome (GRCh38) using minimap2 ^54^. According to the genomic coordinates of the mapped molecules, the structures of each isoform represented in the gtf format were obtained by utilizing the script ‘collapse_isoforms_by_sam.py’ from the cDNA cupcake toolkit ^55^ (v27.0.0, https://github.com/Magdoll/cDNA_Cupcake).

### Isoform quality control, filtering and classification

We merged the isoforms identified across different samples with gffcompare ^56^. We then utilized SQANTI3 (v3.3) ^57^ to compare the identified isoforms against the reference transcriptome annotation, which produced quality control and classification information for each isoform. To provide a comprehensive reference for SQANTI3, we merged databases from multiple sources, including GENCODE (Release v37 for GRCh38) and RefSeq (NCBI Homo sapiens Annotation Release 109 for GRCh38), for the annotation of isoform structures. With this assembled reference, SQANTI3 classified each isoform into FSM, ISM, NIC, NNC and several other structural types ^57^. The rule-based filter functionality of SQANTI3 was used to filter out the isoforms that were likely to be sequencing artifacts such as intra-priming, reverse transcriptase template switching (RTS), and non-canonical splice sites with low read coverage. The remaining isoforms and their associated barcodes are assembled into an isoform-level expression matrix 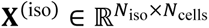, where *N*_iso_ is the number of isoforms, and *N*_cells_ is the number of all cells associated with those isoforms. Finally, we searched the barcodes from PacBio data against the cell barcodes detected in the same sample in the 10x Illumina sequencing data to calculate the expression (UMI counts) of each isoform in each cell. To evaluate the quality of the identified isoforms, we examined whether the 5’ and 3’ ends of each isoform are supported by the TSS and PAS reference. The TSS reference was a union of TSSs from GENCODE, RefSeq and refTSS (v3.1) ^58^, while the PAS reference contains all transcript 3’ ends in GENCODE, RefSeq and PAS from PolyASite (v2.0) ^59^ and PolyADB (v3.2) ^60^. Furthermore, splice junctions from each identified isoform were overlapped with junction reads extracted from the RNA-seq data in The Cancer Genome Atlas colon adenocarcinoma cohort (TCGA-COAD).

### Splice event extraction and analysis

For all isoforms that had passed the SQANTI3’s rule-based filter, we utilized SUPPA (v2.3) ^28^ for the extraction of seven types of AS events including alternative 5’ splice sites (A5), alternative 3’ splice sites (A3), alternative first exon (AF), alternative last exon (AL), skipped exon (SE), mutually exclusive exon (MX) and retained intron (RI). SUPPA analyzed the gtf file obtained from the isoform calling and quality control pipeline to produce seven event files (the ‘ioe’ files) each containing one of the above event types based on the pairwise comparison of the input isoform structures. Taking into consideration the accuracy of splice sites and the inaccuracy of TSS and TTS, we required SUPPA2 to use stringent boundaries for the splice sites of the seven event types, but flexible boundaries (with a variability of 48nt) for the TSS and TTS coordinates of AF and AL.

### Identification of dysregulated gene expression and transcript structures

Dysregulated gene expression (DGE) was identified by comparing the expression of genes (log-normalized UMI) in cells from one group (cell type or subtype) to that from another group. In brief, for each gene, we first counted the number of cells (*n*_1_) expressing this gene with the expression threshold (0) and the total number of cells (*n*_2_) in each group. Next, we applied Fisher’s exact test on the 2 × 2 contingency table formed by *n*_1_ and *n*_2_ from two groups. In addition, we also applied the Mann-Whitney U test (Wilcoxon Rank-Sum test) on the gene expression (log-normalized UMI) in cells from different groups. A significantly upregulated gene was identified by requiring: (1) Benjamini-Hochberg (BH) - adjusted p values (FDR) from Fisher’s exact test < 0.01, (2) odds ratio > 2 and (3) log2 fold change > 0. Significantly downregulated genes were identified using the identical p-value threshold, but with the opposite directional changes (log2 fold change < 0 and odds ratio < 1/2). Dysregulation in isoform expression was identified similarly using the expression of each isoform. For evaluating concordance of DGE quantified by the two sequencing methods, we used genes showing significant changes in both measurements.

To identify dysregulated transcript structures (DTSs), we counted the number of PacBio UMIs supporting a specific isoform versus all alternative isoforms of the same gene for cells from the two groups. The significance of DTSs was computed using Fisher’s exact test on the 2 × 2 contingency table formed by four numbers from the two groups of cells. The significant DTSs were identified by requiring: (1) BH-adjusted p-value (FDR) < 0.01, (2) the percentage of isoform changes > 0 and (3) odds ratio > 2, or (1) BH-adjusted p-value (FDR) < 0.01, (2) the percentage of isoform changes < 0 and (3) odds ratio < 1/2.

### GO enrichment analysis

The list of genes with DTS from comparing EpiT to EpiN was uploaded to WebGestalt ^61^ as input and all the genes with detected expression in these two cell types were used as the background to identify the enriched GO terms (FDR < 0.05, enrichment score > 2).

### Calling RNA editing events

We assembled a comprehensive reference of potential RNA editing sites from three different sources: (1) the RADAR database ^62^, (2) the DARNED database ^63^ and (3) Tan et al., 2017. ^64^. For each putative RNA editing site in the reference, we systematically examined the evidence of reads supporting an A>G change at that locus (or a T>C change on the negative strand) using the ‘mpileup’ subcommand of bcftools ^65^. We then associated each event with the isoforms and cell barcodes from which the event was discovered. To remove interference from genomic SNVs, we searched the candidate RNA editing sites against the dbSNP database and discarded the sites with an allele frequency greater than 0.01 in any genomic studies it had been concerned. We retained RNA editing sites with more than 10 read coverage and computed an edit ratio for each site, defined as the number of edited reads divided by the number of total read coverage. For these RNA editing sites, we additionally calculated their per isoform edit ratio using the reads that can be unambiguously identified to a specific isoform and if the per isoform read coverage is greater than five. The per isoform edit ratio is defined as the number of edited reads divided by the number of total read coverage from the isoform (**Fig. 4a**).

### Lineage and trajectory analysis

We employed monocle 3 ^36^ for the single-cell lineage and trajectory analysis on the epithelial component of the short-read scRNA-seq dataset. The principal graph was constructed using the ‘learn_graph’ subroutine of monocle 3 with default parameters using the tSNE coordinates and the nodes corresponding to the four stem cells subclusters were designated as the root for diffusion pseudotime inference. The epithelial cells that were also detected in the PacBio long-read scRNA-seq were associated with their isoform level expression values. The expression of genes and isoforms, as well as the percentage of isoforms that were associated with a specific lineage, were identified based on Moran’s I statistic ^66^.

### Quantification of isoforms in TCGA samples

Isoform-level expression of the TCGA-COAD bulk samples was quantified using stringtie (v2.2.1) ^67^ using the merged transcriptomic reference of novel identified isoforms, GENCODE (version 37) and RefSeq (NCBI Homo sapiens Annotation Release 109 for GRCh38).

### Estimation of the signature scores of EpiT subtypes in bulk samples

We identified the top differentially expressed genes in the EpiT subtypes cE02 and cE03 (**Supplementary Table 6**). We then computed the z-scores of those up- and down-regulated genes in each bulk sample. The signature score of an EpiT subtype is defined as the averaged z-scores of the up-regulated genes minus the averaged z-scores of the down-regulated genes.

### Survival analysis

We correlated the gene expression or activity scores with the PFS and OS of the CRC patients based on the clinical data from TCGA-COAD and CRCSC using the Cox proportional hazards regression model. The Kaplan-Meier method was used to estimate the survival function of the patients under each condition.

### Proteomic analysis of isoforms

To comprehensively characterize the proteome in CRC, GeneMarkS-T ^68^ was used to predict ORFs in the PacBio-identified isoforms. For comparison and annotation of the predicted ORFs, we collected the human reference proteome from Uniprot (release 2022-02) ^69^, including all sequences from the reviewed (SwissProt), unreviewed (TrEmbl) and spliced isoform (Varsplice) collections at protein existence (PE) level PE=1 and PE=2. Using blastp ^70^, we searched each predicted ORF sequence against the UniProt reference proteome, and the reference sequence with the highest alignment score was assigned to it as its nearest homolog in the database. The ORFs with a similarity of less than 99% to their nearest homolog in the reference were considered novel protein sequences. Protein domains of each ORF sequence, as defined by the Pfam-A hidden Markov model (HMM)-based multiple sequence alignments ^71^, were annotated by hmmscan (http://hmmer.org/). We regarded a novel ORF sequence as having a domain gain if it contained a proper superset of the Pfam domains from its nearest Uniprot homolog, or a domain loss if it contained only a proper subset of them. We also inferred the sensitivity of the PacBio-identified isoforms to nonsense-mediated decay (NMD) based on the relative position of the predicted stop codon from the last splice junction. Lastly, the subcellular localization of the ORF-derived proteins was predicted using DeepLoc ^72^.

### Mass spectrometry data analysis

To validate the predicted ORFs, particularly the novel ORFs, we collected mass spectrometry (MS) data from three independent sources: (1) a CRC cohort from the National Cancer Institute’s Clinical Proteomic Tumor Analysis Consortium (CPTAC), (2) an in-house MS dataset consisting of 22 samples from 11 CRC patients (11 tumor and 11 paired adjacent normal), and (3) an in-house MS data from a colon cancer cell line, HCT116. With the same pipeline as a previous study ^73^, we searched MS2 spectra from the MS data against a reference by integrating annotated protein sequence from UniProt (release 2022-02) and novel ORFs predicted based on isoforms derived from LR-seq using Comet ^74^. Five peptides were reported for each spectrum query and these peptides were further ranked and scored for final identification using Percolator ^75^. The validated novel ORFs were defined by its inclusion of at least one novel peptide (FDR < 0.01) in the MS data that does not match with any protein sequence from the UniProt database.

### Proposing putative neoepitopes for cancer vaccine development

To obtain a shortlist of tumor-specific novel transcript isoforms, we extracted the NIC and NNC isoforms identified in our long-read scRNA-seq profiles and quantified their expression levels in the TCGA-COAD bulk samples along with the known isoforms in GENCODE and RefSeq. Tumor-specificity was ensured by both filtering for isoforms discovered only from the EpiT cells and not expressed (TPM < 0.5) in the TCGA-COAD normal samples. Additionally, we required each novel isoform to possess at least one novel junction which has supporting reads in <5% of TCGA normal samples and >5% TCGA tumor samples. To ensure active translation of the isoforms, we prioritized 12 isoforms from the shortlist whose ORF subsequences were supported at least once by the MS data from tumor samples but without any peptides detected in MS data from normal samples. Out of five isoforms in this shortlist selected for experimental confirmation, we validated the unique splice junction sequences and RT-PCR products for three isoforms arising from AF and A5SS AS events, and one isoform with retained introns (RI) in CRC cell lines, as well as at least two of the five matched patient samples tested.

To derive the set of tumor-specific neoepitopes from the tumor-specific novel transcript isoforms, we assembled a collection of all the *k*-mers (*k* = 9) that are present in the predicted ORF sequences and further curated a subset containing tumor-specific novel *k*-mers by requiring: (1) the *k*-mers are not in the Uniprot reference, (2) the *k*-mers are only present in ORFs in transcript isoforms from EpiT but not from EpiN or other cell types, (3) the *k*-mers are from the transcript isoforms that are expressed (TPM > 0.5) in more than five TCGA-COAD tumor samples and not expressed (TPM < 0.5) in the normal samples (4) the *k*-mers are from the ORFs in the transcript isoforms classified as NIC and NNC. In this way, we obtained a set *S* of candidate neoepitope *k*-mers that are from recurrent tumor-specific isoforms across tumor samples.

For a given population, we predicted the binding affinity of each neoepitope *k*-mer against the frequent HLA alleles in the population (represented as the set ℋ) using netMHCpan (v4.1) ^76^. We used the rank of elution score, EL_rank < 2 as the threshold of binding/non-binding for a *k*-mer to a specific HLA allele. In this way, we obtained a mapping *M* that represented the HLA alleles that a neoepitope *k*-mer can bind to. For example, ℳ[*s*] (ℳ[*s*] ⊆ ℋ) contains the HLA alleles to which neoepitope s (*s* ∈ *S*) binds.

To optimize a list of neoepitope *k*-mers with a high probability of containing at least one epitope with binding potential to a patient’s HLA allele for a given population, we searched the combination of neoepitope *k*-mers in the shortlist to cover as many frequent HLA alleles in a population as possible. Such an optimization task was formulated as a maximum coverage problem:

Input: The size of the shortlist n and the sets of HLA alleles covered by each epitope {*M*[*s*_1_], *M*[*s*_2_], …, *M*[*s*_|*S*|_]}.

Output: The subset *S*′ ⊆ *S* with |*S*′| ≤ *n* such that ⋃_{*s*∈*S*′}_ *M*[*s*]| is maximized.

Additionally, we took into account the HLA’s allele frequency by assigning it as the weight in a weighted maximum coverage problem:

Input: The size of the shortlist n, the sets of HLA alleles covered by each epitope {*M*[*s*_1_], *M*[*s*_2_], …, *M*[*s*_|*S*|_]}, and the weight of each HLA allele *a* (*a* ∈ ℋ), *W*[*a*].

Output: The subset *S*^′^ ⊆ *S* with |*S*′| ≤ *n* such that 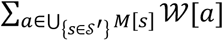 is maximized.

As both the weighted and unweighted maximum coverage problems are NP-hard ^77^, we used the following greedy algorithm for the selection of an epitope shortlist with size *n* (**Algorithm 1**, S_ELECT_-K_MERS_-F_OR_-P_OPULATION_), **which is also illustrated in Extended Data Fig. 11A**. We omitted the algorithm for the unweighted version in the presentation here as it is a special case of the weighted version when *W*[*a*] = 1 (∀*a* ∈ ℋ). On each iteration, we selected an epitope *k*-mer not yet in the shortlist that would result in the maximum gain in the weight of covered HLA alleles (**Algorithm 1**, S_ELECT_-B_EST_-K_MER_). If the HLA alleles were already completely covered before the size of the shortlist reaches n, we reset the set of uncovered HLA alleles and continued the iteration. In this way, the remaining epitopes in the shortlist were optimized again for the maximum coverage of HLA alleles in the population.

To optimize the neoepitope shortlist for the TCGA-COAD patients, we first obtained the HLA subtyping of these patients from Thorsson et al. ^78^ and computed the binding affinity of the neoepitopes in set *S* to all the HLA alleles found in them. We defined the coverage of a neoepitope for a patient as at least one HLA allele of the patient covered by the neoepitope. We then optimized the shortlist of neoepitopes for the maximum coverage of TCGA patients by solving a similarly-defined unweighted maximum coverage problem for the population HLA alleles. This resulted in a similar procedure (**Algorithm 1**, S_ELECT_-K_MERS_-F_OR_-P_ATIENT_-G_ROUP_) as above, by redefining *M*[*s*] to be the patients covered by neoepitope *s*.

#### Algorithm 1 Select Splicing-derived Neoepitopes for Cancer Vaccine Development

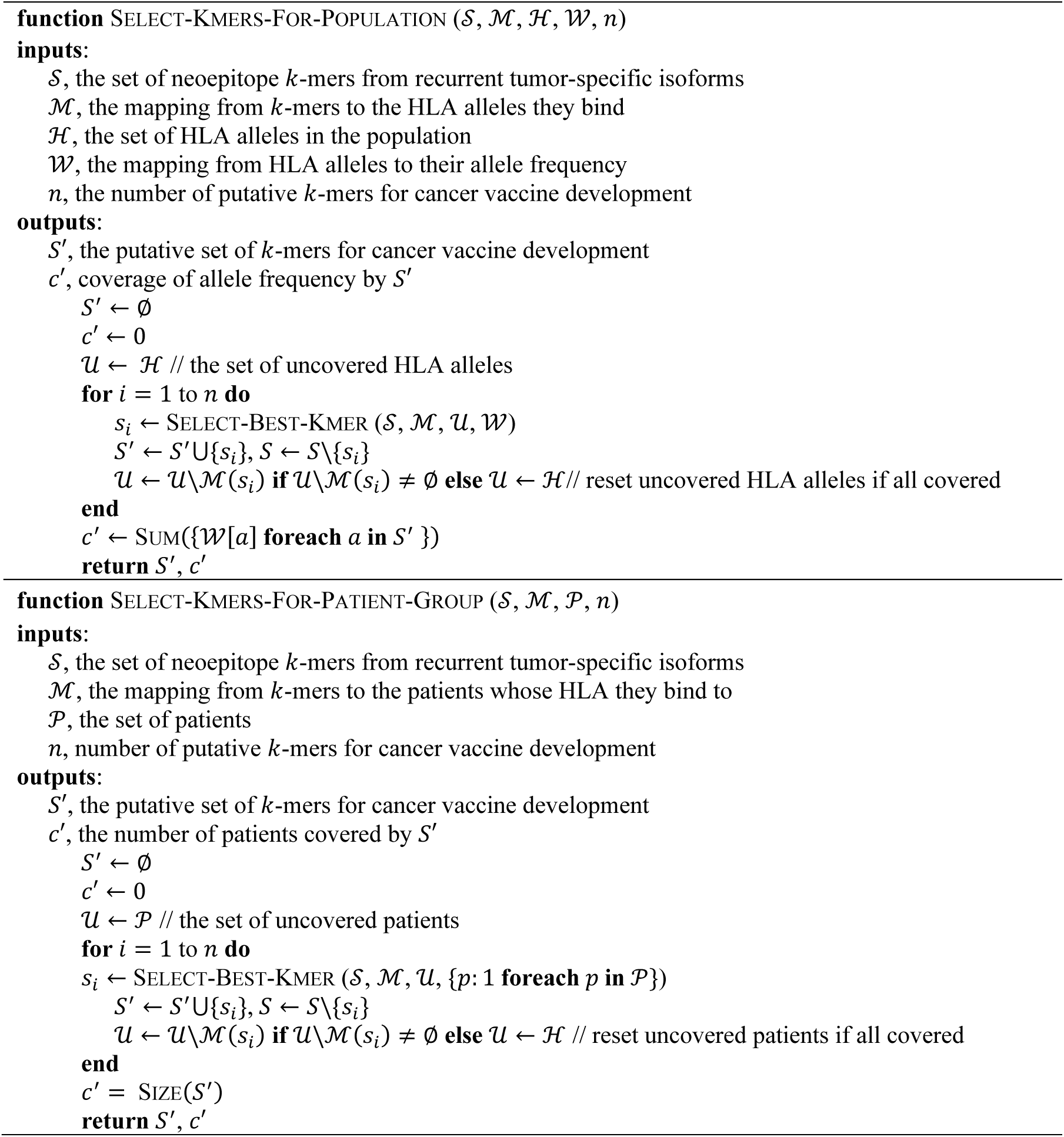
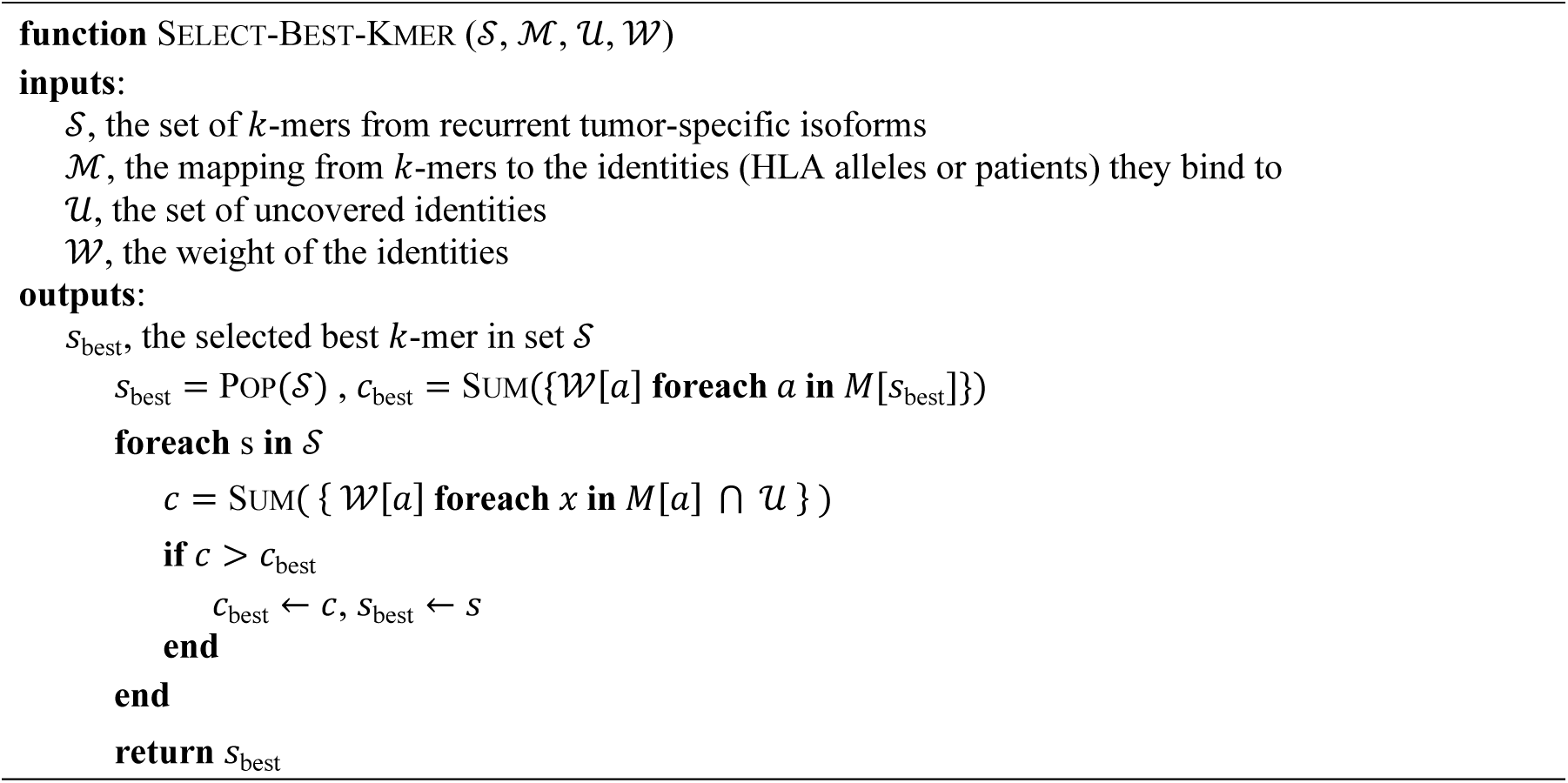

## Acknowledgments

We thank all past and present YT and XG lab members for their constructive feedback on this project, Ng Desi for assisting with the single cell isolation procedure, and Luke Esau and other members from bioscience core laboratory at KAUST for performing the single-cell Illumina and PacBio sequencing. We additionally thank personnel from Iain Bee Huat Tan’s lab for providing the iCMS classification on the c295 dataset. The results in the study are in whole or part based upon data generated by the TCGA Research Network: https://www.cancer.gov/tcga.

This study is supported by the National Research Foundation Singapore and the Singapore Ministry of Education under its Research Centres of Excellence initiative. Y.T. is funded by NMRC OF-IRGs (NMRC/OFIRG/MOH-000380, MOH-000923). X.G. is supported by the King Abdullah University of Science and Technology (KAUST) Office of Research Administration (ORA) under Award No FCC/1/1976-44-01, FCC/1/1976-45-01 and REI/1/5234-01-01.

## Authors’ Contributions

Z.L. and B.Z. performed computational analysis. J.J.C. performed experiments, single cell isolation and library preparations, as well as analyzed data. H.T., Q.Y.T. and X.H.C. performed single cell isolation, RNA extractions and tissue sample preparation for mass spectrometry. P.D. constructed PacBio libraries and performed sequencing. X.F. assisted with computational analysis. C.C. and D.K. performed the mass spectrometry acquisition. F.C. assisted in mass spectrometry sample preparation. S.W., B.E.S., I.J.W.T., K.Y.L., B. L., W.K.C. and K.K.T provided COAD clinical samples. Z.L, B.Z., J.J.C., X.G. and Y.T. designed the study and prepared the manuscript. All authors reviewed and commented on the manuscript.

## Data and Code Availability

Raw sequencing data used in this study are deposited in European Nucleotide Archive under session number PRJEB61296. The identified RNA transcript isoforms from LR-seq data are visualized and available for downloading from UCSC genome browser tracks with the link: https://genome.ucsc.edu/s/binzhang/COAD_Colored_PacBio. Code for data analysis and neoepitope selection is available at https://github.com/lzx325/CRC-atlas.git.

