## Supplementary Notes for "An isoform-resolution transcriptomic atlas of colorectal cancer from long-read single-cell sequencing"

#### Section S1 The proteomic analysis of isoforms in the long-read transcriptomic atlas

To investigate the effect of transcript structure alterations on the proteome, we predicted the open reading frames (ORFs) of all the identified isoforms using GeneMarkS-T (Tang et al., 2015). The majority of the isoforms were predicted to have an ORF, with FSM having the highest percentage (92.56%) followed by ISM (84.75%), NIC (85.77%) and NNC (78.62%) (**Extended Data Fig. 9a**). Almost all predicted ORFs started with the canonical start codon 'AUG' and most (>85%) from the FSM, NIC and NNC categories terminated with a canonical stop codon (**Extended Data Fig. 9b and 9c**). However, only 48.56% of the ISM isoforms have a canonical stop codon, suggesting that many of them were are only fragments of the original transcript (**Extended Data Fig. 9c**). Compared to FSM and ISM, the novel splice junctions of NIC and NNC isoforms resulted in a higher potential of being targeted by the nonsense-mediated decay (NMD) pathway (**Extended Data Fig. 9d**). To identify potential novel peptides sequences, we searched the ORF sequences against the reference human proteomic sequences in Uniprot (release 2022-02) (**Methods**). Over 82% and 88% of the ORF sequences from the FSM and ISM isoforms, respectively, had >99% similarity matches in Uniprot. Over 34% of the ORF sequences from NIC and NNC are novel predictions, highlighting the contribution of AS to the proteomic diversity (**Extended Data Fig. 9e**).

#### Section S2 Command lines of software tools used in this study

<u>*Mapping in-house short-read scRNA-seq profiles to bc295:*</u>

```
anchors <- FindTransferAnchors(
    reference = [reference Seurat object],
    query = [query Seurat object],
    dims = 1:30
)
prediction_cluster_midway = TransferData(
    anchorset = anchors, refdata = [reference Seurat object 'ClusterMidway' labels],
    dims = 1:30
 )
```

<u>*Alignment of the long-read molecules to genome using minimap2:*</u>

```
minimap2 -t 30 -ax splice -uf --secondary=no -C5 [path to genome fasta file]  [PacBio molecule fasta file] > [alignment output file]
```

<u>*Identification of long-read isoforms using cDNA cupcake:*</u>

```
collapse_isoforms_by_sam.py --input [PacBio molecule fasta file]  \
```

```
    --bam [alignment output file] -c 0.99 -i 0.95 \
    --gen_mol_count \
    -o [output file prefix] \
    --cpus 20
```

*Classification of isoforms using SQANTI3:*

```
sqanti3_qc.py \
   --gtf [path to cDNA cupcake's gff output file] \
   [path to the reference transcript isoform gtf file] [path to the genome fasta file] \
   --fl_count [path to cDNA cupcake's isoform abundance file] \
   --cage_peak [reference CAGE peaks file] \
   --polyA_motif_list [reference polyA motifs] \
   --polyA_peak [reference polyA peaks file] \
   --dir  [output directory]
```

*Subtyping of the normal and tumor epithelial cells using xgboost:*

Training the xgboost model:

```
library(xgboost)
bst <- xgb.train(
    data = dtrain,
    max.depth = 4, eta = 0.5, nthread = 50, nrounds = 200, objective = "multi:softmax",
    eval_metric="merror",num_class=n_class,
    verbose=2)
)
```

Inference with the trained xgboost model:

```
Prediction <- predict(bst,dtest)
```

*Genotyping of patients' HLA alleles using arcasHLA:*

```
arcasHLA extract --single [path to the bam file of long-read or short-read sequencing] -o [output directory]
arcasHLA genotype --single --min_count 3 [arcasHLA extracted fastq file] -g A,B,C -o [output directory] -t 8
```

*Prediction of the HLA binding affinity of neoepitopes using netMHCpan:*

```
netMHCpan -a [MHC allele] -l 9 -f [neoepitope fasta file]
```

*Peptides identification from MS data with Comet and Percolator:*

```
comet.linux.exe -Pcomet_param_file [path to the mzML file]
```

crux percolator --overwrite T --output-dir sample --test-fdr 0.1 --train-fdr 0.1 [path to the pin format file outputted by comet]

*<u>Some parameters in comet_param_file was shown below:</u>*

decoy_search = 2

peptide_mass_tolerance = 3

peptide_mass_units = 2

mass_type_parent = 1

mass_type_fragment = 1

precursor_tolerance_type = 0

isotope_error = 0

search_enzyme_number = 1

num_enzyme_termini = 2

allowed_missed_cleavage = 2

variable_mod01 = 15.9949 M 0 3 -1 0 0 0.0

max_variable_mods_in_peptide = 5

require_variable_mod = 0

fragment_bin_tol = 1.0005

fragment_bin_offset = 0.4

theoretical_fragment_ions = 1

output_percolatorfile = 1

ms_level = 2

sample_enzyme_number = 1

digest_mass_range = 600.0 5000.0

peptide_length_range = 5 63

max_duplicate_proteins = 20

max_fragment_charge = 3

max_precursor_charge = 6

minimum_peaks = 10

remove_precursor_tolerance = 1.5

### Extended Data Figures

**Extended Data Figure 1 Preprocessing of the Illumina short-read scRNA-seq data. Related to Fig. 1.**

(a) The Illumina short-read scRNA-seq data of the normal samples were mapped to the genome using 'cellranger count'. 'cellranger aggr' was used to perform cross-library normalization. We separately integrated the normal and tumor samples using the Seurat subroutine 'IntegrateData'. Finally, the integrated normal and tumor samples were mapped (using the 'TransferData' subroutine) to the c295 dataset for cell type identification.

(b) Distribution of the number of genes, UMIs, and percentage of mitochondrial expression of the short-read scRNA-seq data. The cutoffs for the three metrics are shown respectively.

(c) Distribution of the number of cells per sample of the short-read scRNA-seq data. We achieved slightly lower but comparable cell count as compared to related single-cell studies using similar technologies.

(d) Expression of epithelial cell markers among identified epithelial cell types.

(e) Expression of cell markers among identified myeloid and fibroblast cell type(s).

(f) Expression of cell markers among identified lymphoid cell types.

**Extended Data Figure 2 Statistics of transcript isoform and cell detection of PacBio long-read scRNA-seq data. Related to Fig. 2.**

(a) Percentage of transcript isoforms in each structural category.

(b) Statistics of normal-only, tumor-only and common transcript isoforms by each structural category.

(c) Overlap of cell barcodes between short-read and long-read sequencing in each sample.

**Extended Data Figure 3 Correlation of long-read and short-read scRNA-seq among the commonly detected cells. Related to Fig. 2.**

(a) Statistics of cell types among the common cells detected by long-read and short-read sequencing.

(b) Number of commonly detected genes by long-read and short-read sequencing per cell type.

(c) Scatterplot (each dot represents a gene) showing the correlation of gene expression levels by long-read and short-read sequencing within EpiT and EpiN.

**Extended Data Figure 4 Dysregulated transcript structures and the definition and statistics of the SUPPA2 events. Related to Fig. 3.**

(a) The number of significantly up- and downregulated isoforms and their overlap with isoforms with increased, decreased and not significantly changed (NS) percentages in EpiT compared to EpiN.

(b) Schematic showing the definition of the seven types of SUPPA2 events.

(c) Statistics of the known and novel SUPPA2 events in each category.

**Extended Data Figure 5 Statistics and validation of RNA editing events. Related to Fig. 4.**

(a) Number of RNA editing events in coding regions, 5'UTRs, 3'UTRs and non-coding transcripts.

(b and c) Validation of RNA editing events by Sanger sequencing in (B) a colon epithelial cell line, CCD 841 CoN, and two CRC cell lines, DLD-1 and HCT116, and (C) five in-house matched patient samples.

(d) REPIs of the identified RNA events with sufficient per isoform sequencing read support. The events that are shown in **Fig. 4b** are annotated.

**Extended Data Figure 6 Differentially expressed genes in the EpiN subtypes. Related to Fig 5.**

(a) Volcano plot showing the top differentially expressed genes in the four comparisons between EpiN subtypes: (1) Stem vs. rest, (2) enterocyte lineage vs. stem, (3) goblet lineage vs. stem, (4) *BEST4* lineage vs. stem.

(b) Number of significantly up- and downregulated genes in the stem, goblet, enterocyte and *BEST4* subtypes.

(c) Overlap of differentially expressed genes in the above four comparisons.

**Extended Data Figure 7 Differentially expressed genes in the EpiT subtypes. Related to Fig. 6.**

(a) Distribution of iCMS2 and iCMS3 epithelial cells in the in-house dataset.

(b) Distribution of iCMS2 and iCMS3 epithelial cells, and the three EpiT subtypes in the c295 dataset.

(c) Volcano plot showing the top differentially expressed genes in the four comparisons between the three EpiT subtypes and their corresponding subtypes in EpiN (cE01, cE02, cE03).

(d) Overlap of common differentially expressed genes in the above four comparisons.

**Extended Data Figure 8 Association of patient survival with the EpiT subtypes. Related to Fig. 6.**

(a) Proportion of upregulated (up), downregulated (down) and not significantly changed (NS) genes for genes with DTS, without DTS and those with only one detected isoform in each EpiT subtype.

(b) Overlap of common genes with DTS in the three EpiT subpopulations.

(c) Overall survival (OS) of TCGA-COAD patients with different expression levels of *TFF1*.

(d)The number of patients with different combinations of cE02 and cE03 signature scores. H: high, L: low.

(e) OS of TCGA-COAD patients with different activities of cE02 and cE03 signature genes.

(f) OS and (g) progression-free survival (PFS) of patients in the CRCSC microarray data with different cE02 and cE03 signature scores.

**Extended Data Figure 9 Statistics of the predicted proteome of the patient samples using the long-read isoforms. Related to Fig. 7.**

(a-d) Percentage of isoform ORFs (a) by each structural category, with a proper (b) start codon and (c) stop codon and (d) that will be potentially targeted by NMD.

(e) Percentage of coding isoforms with a known/novel ORF sequence.

(f and g) Number of coding isoforms with novel ORF sequences that have a hit in three MS datasets: (1) CPTAC COAD, (2) in-house CRC primary tumor and (3) in-house HCT116 cell line, by (f) structural category and (g) the cell type in which detected.

**Extended Data Figure 10 Validation of the unique splice junctions for selected recurrent tumor-specific isoforms. Related to Fig. 7.**

(a) Sanger sequencing validation of splice junctions in normal colon epithelial cells, CCD 841 CoN and a CRC cell line, DLD-1. Novel splice junctions are marked by black dotted lines and the 5' junction of sequences unique to selected isoforms by red dotted lines.

(b) Consensus sequence around splice junctions or unique regions (demarcated by the blue highlighted text) derived from Sanger sequencing of five in-house patient samples.

(c) Validation of the protein overexpression of HA-tagged open reading frames derived from the validated neoepitopes in (b).

(d) Effect of overexpressing the open reading frames derived from the validated neoepitopes on anchorage-independent growth in CCD 841 CoN cells.

**Extended Data Figure 11 Analysis of an alternative neoepitope panel optimized for the global population. Related to Fig. 7.**

(a) Illustration of the neoepitope optimization process. The algorithm is an iterative process. On each iteration, the neoepitope with the maximum coverage of HLA alleles/patients is selected.

(b) HLA binding profile of the 22 neoepitopes from the same 4 isoforms as in **Fig. 7** but optimized for the global population.

(c) Distribution of the per-HLA allele binding affinity of the 22 neoepitopes in **Extended Data Fig. 11b**.

**Extended Data Figure 12 Summary of the bioinformatic analysis pipeline in this study.**

### Supplementary Tables

**Table S1 Sample and library information.**

(a) Patient metadata, demographics, and consensus molecular subtypes (CMS).

(b) Illumina short-read library information.

(c) Comparison of short-read data cell type statistics with c295.

(d) Short-read data cell type statistics by sample.

(e) Statistics of EpiT identification based on SCNV and xgboost.

(f) PacBio long-read library information.

**Table S2 REPS and REPI of identified RNA editing events.**

**Table S3 Differentially expressed genes and dysregulated transcript isoforms in the EpiN subtypes.**

**Table S4 Differentially expressed genes and dysregulated transcript isoforms in the EpiT subtypes.**

**Table S5 EpiT marker genes to compute the signature scores of EpiT subtypes using bulk RNA-seq and microarray data.**

**Table S6 Tumor recurrent novel isoform candidates for neoepitope derivation.**

**Table S7 Proposed panels of neoepitopes for cancer vaccine development.**

**Table S8 HLA genotypes of in-house patients by arcasHLA.**

**Table S9 PCR primer sequences and antibodies used for experimental validation of DGEs, RNA editing events, tumor-specific isoforms deriving the neoepitope panel and their ORFs.**
