## Extended Data Figures for "An isoform-resolution transcriptomic atlas of colorectal cancer from long-read single-cell sequencing"

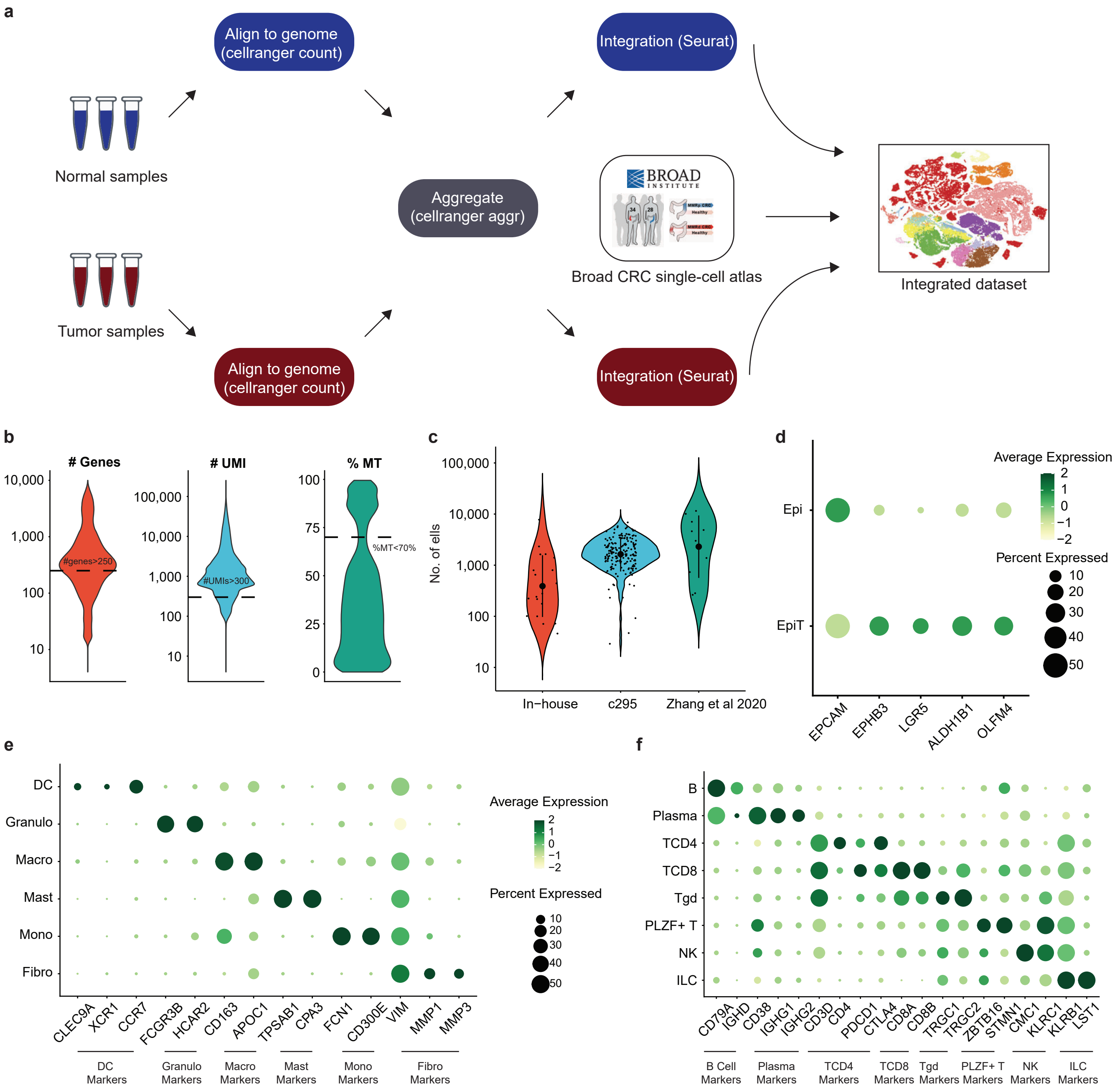


Extended Data Figure 1

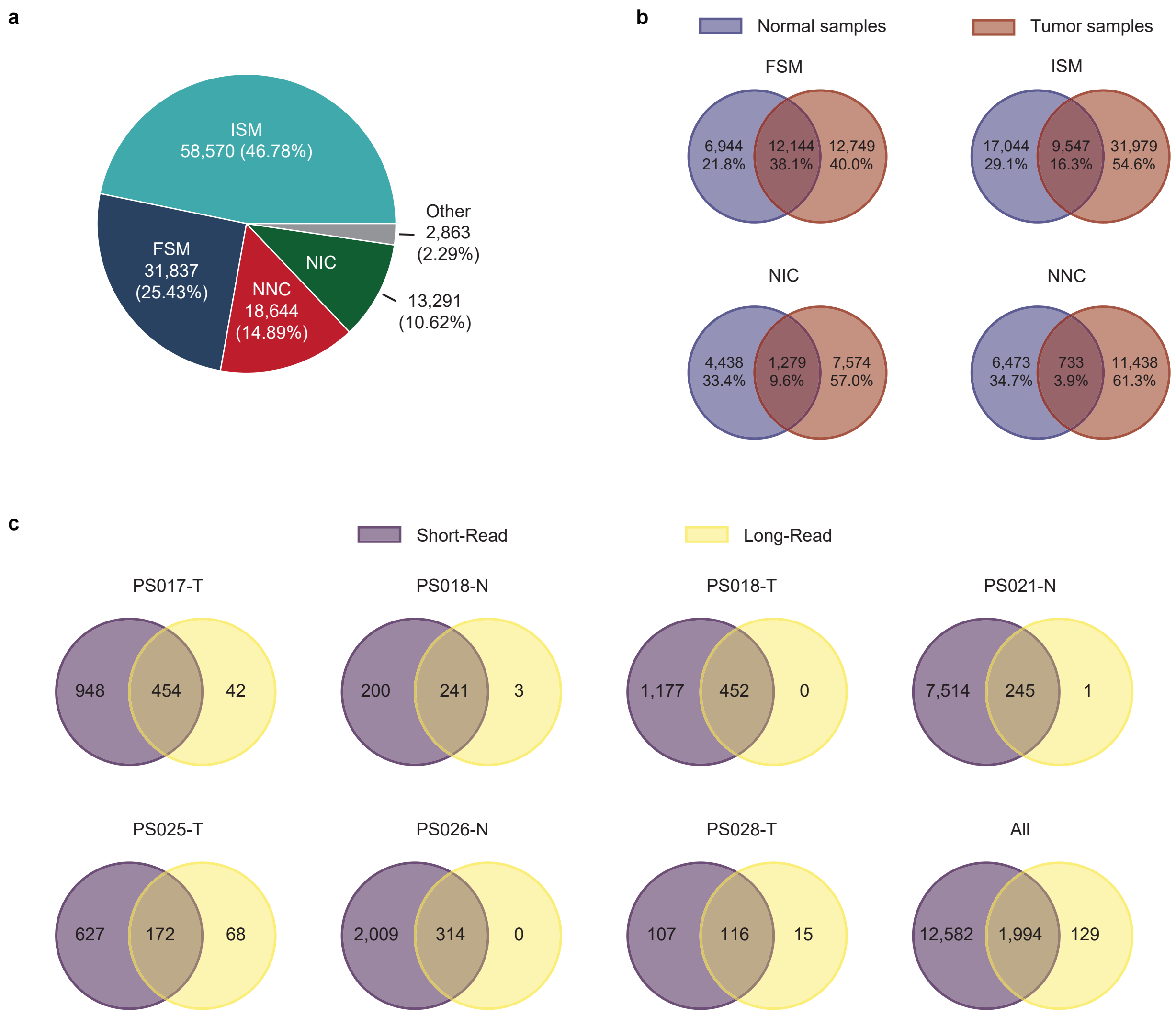
a
ISM
58,570 (46.78%)
FSM
31,837
(25.43%)
NNC
18,644
(14.89%)
NIC
13,291
(10.62%)
Other
2,863
(2.29%)
b
Normal samples
Tumor samples
FSM
6,944 21.8%
12,144 38.1%
12,749 40.0%
ISM
17,044 29.1%
9,547 16.3%
31,979 54.6%
NIC
4,438 33.4%
1,279 9.6%
7,574 57.0%
NNC
6,473 34.7%
733 3.9%
11,438 61.3%
c
Short-Read
Long-Read
PS017-T
948
454
42
PS018-N
200
241
3
PS018-T
1,177
452
0
PS021-N
7,514
245
1
PS025-T
627
172
68
PS026-N
2,009
314
0
PS028-T
107
116
15
All
12,582
1,994
129

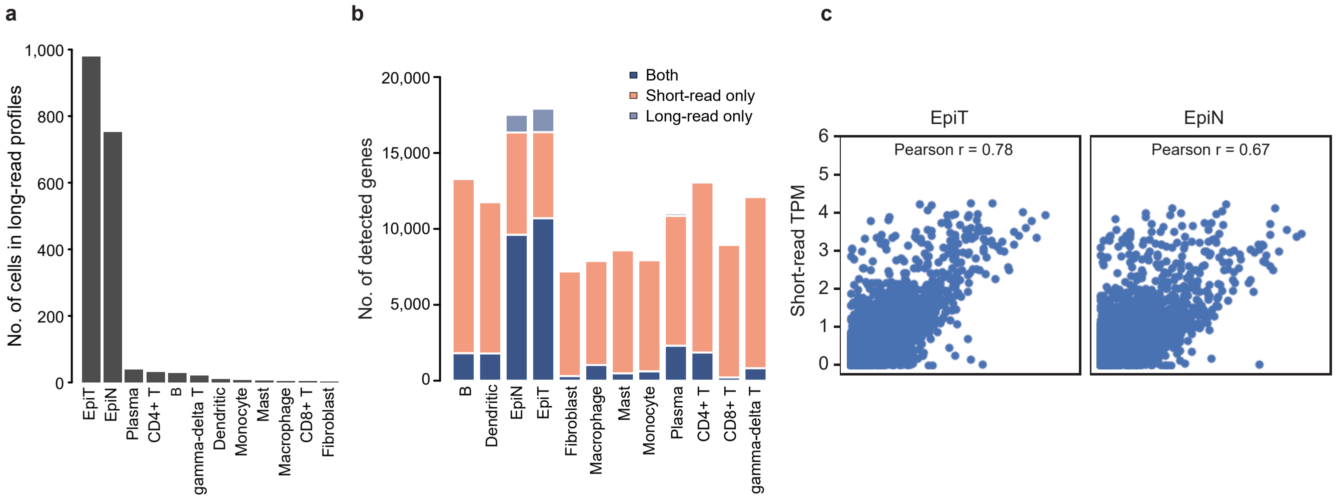
a
No. of cells in long-read profiles
1,000
800
600
400
200
0
EpiT
EpiN
Plasma
CD4+ T
B
gamma-delta T
Dendritic
Monocyte
Mast
Macrophage
CD8+ T
Fibroblast
b
No. of detected genes
20,000
15,000
10,000
5,000
0
Both
Short-read only
Long-read only
B
Dendritic
EpiN
EpiT
Fibroblast
Macrophage
Mast
Monocyte
Plasma
CD4+ T
CD8+ T
gamma-delta T
c
Short-read TPM
EpiT
EpiN
Pearson r = 0.78
Pearson r = 0.67
6
5
4
3
2
1
0

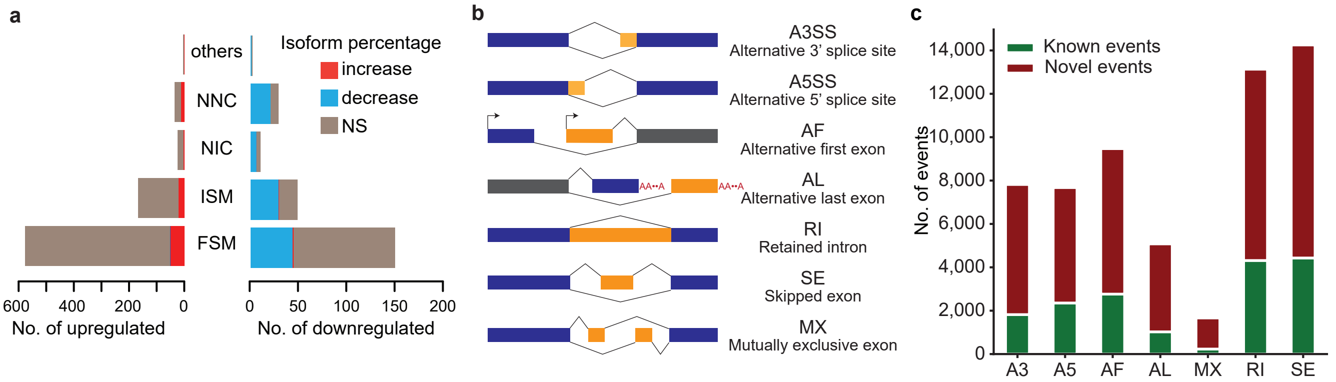


Extended Data Figure 4

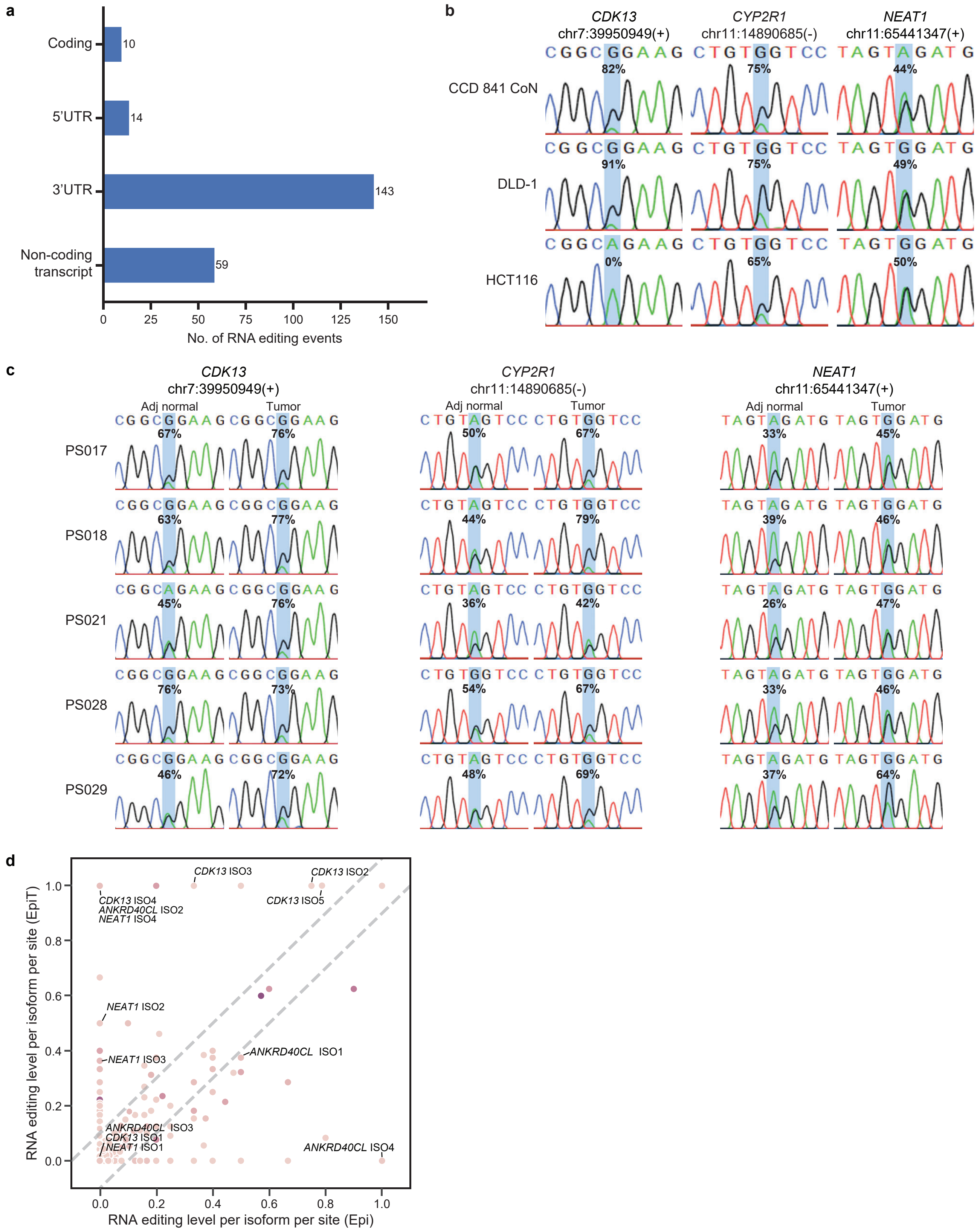
a
Coding
5'UTR
3'UTR
Non-coding transcript
10
14
143
59
No. of RNA editing events
b
CDK13 chr7:39950949(+)
CYP2R1 chr11:14890685(-)
NEAT1 chr11:65441347(+)
CCD 841 CoN
DLD-1
HCT116
c
Adj normal
Tumor
PS017
PS018
PS021
PS028
PS029
d
RNA editing level per isoform per site (EpiT)
RNA editing level per isoform per site (Epi)
CDK13 ISO3
CDK13 ISO2
CDK13 ISO5
CDK13 ISO4
ANKRD40CL ISO2
NEAT1 ISO4
NEAT1 ISO2
NEAT1 ISO3
ANKRD40CL ISO1
ANKRD40CL ISO3
CDK13 ISO1
NEAT1 ISO1
ANKRD40CL ISO4

Extended Data Figure 6

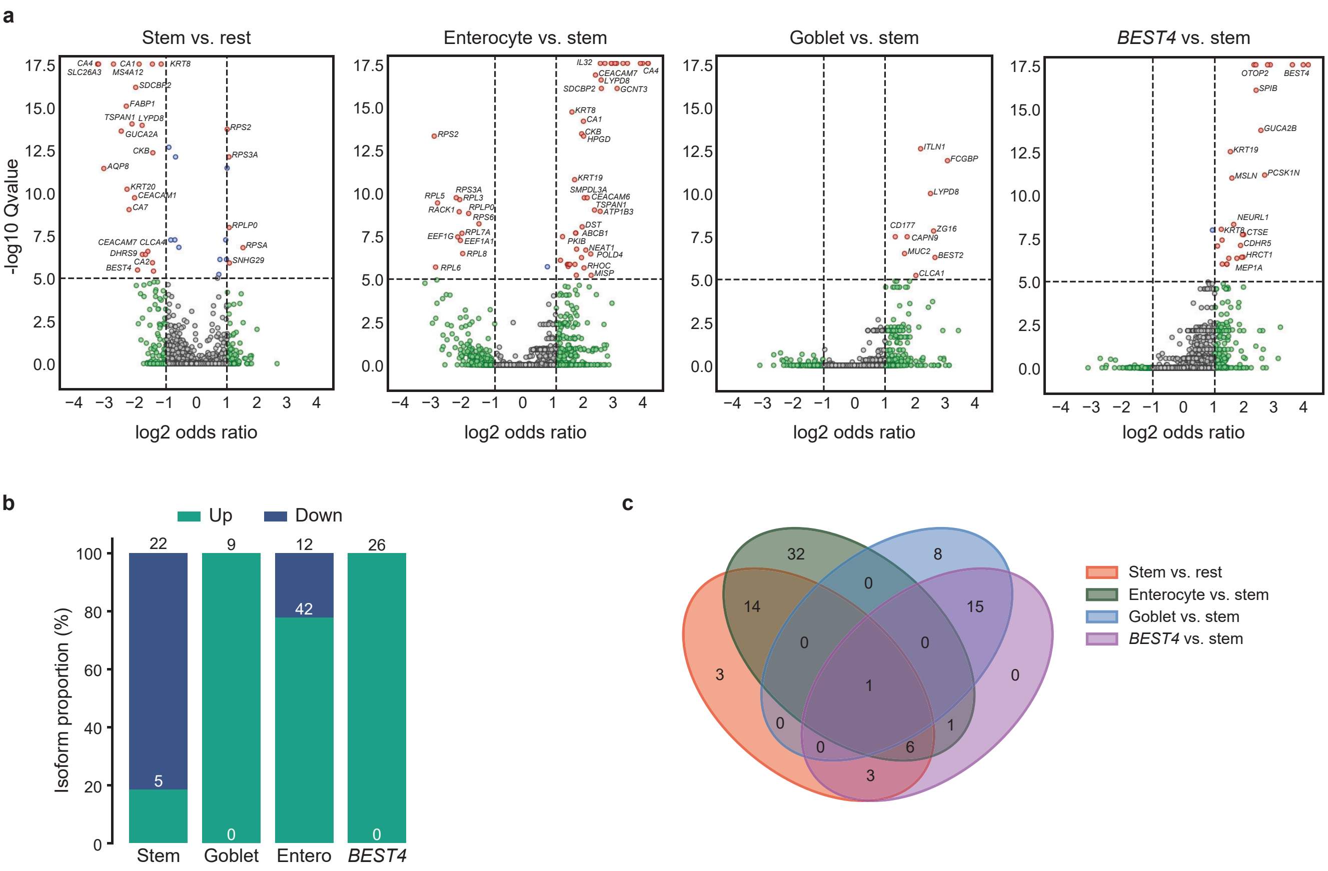

Extended Data Figure 7

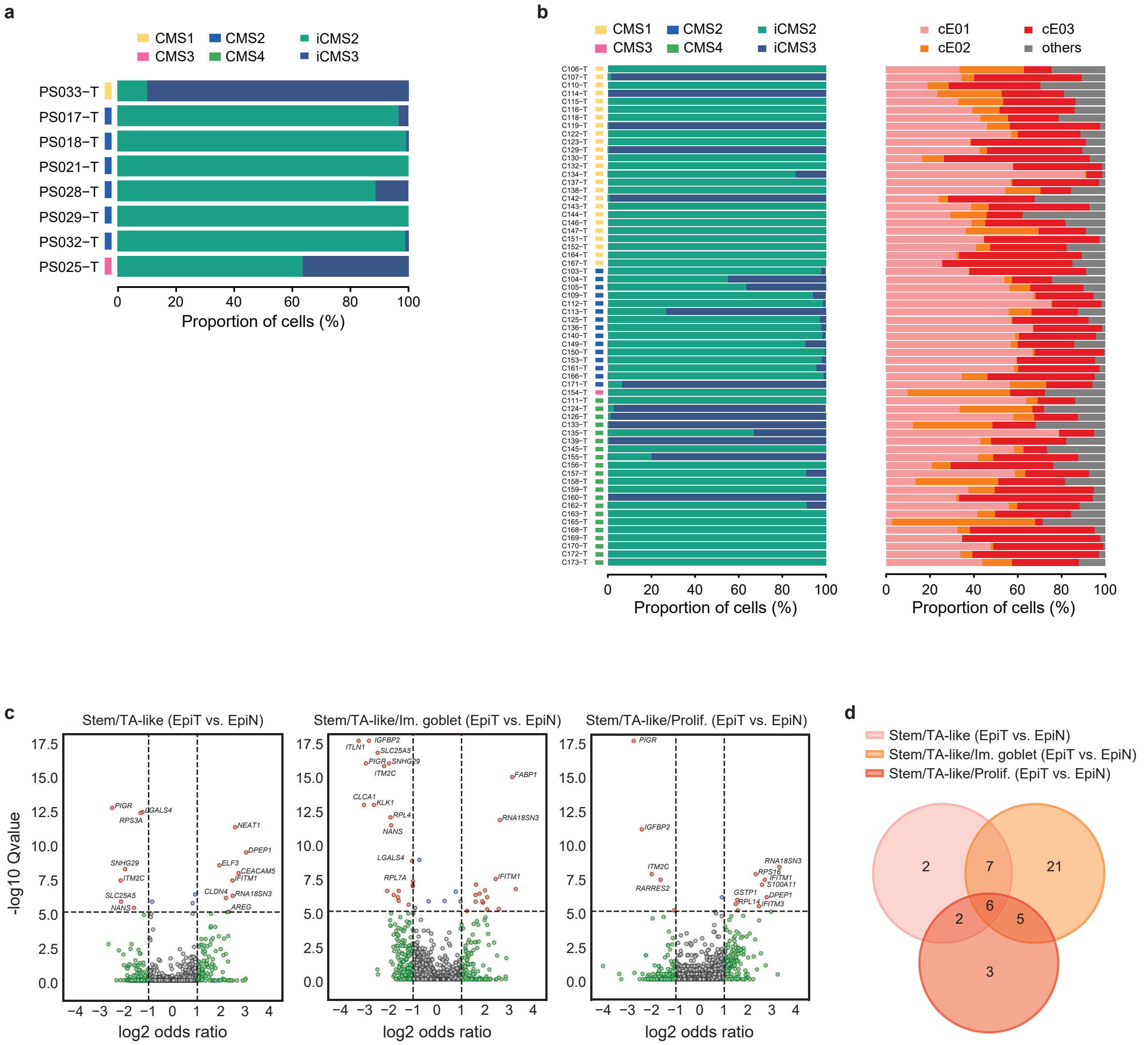


Extended Data Figure 7

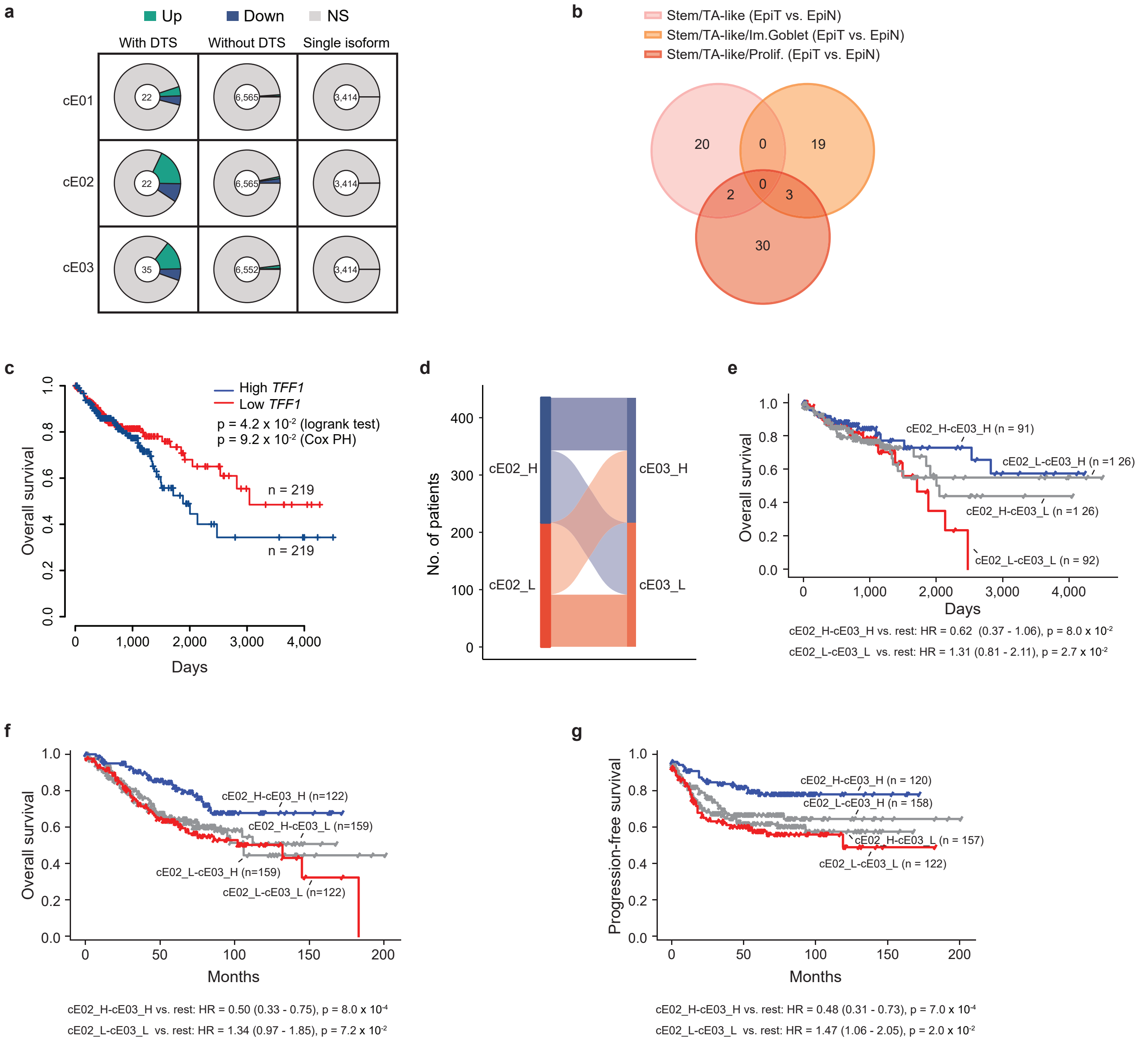
a
Up
Down
NS
With DTS
Without DTS
Single isoform
cE01
cE02
cE03
22
6,565
3,414
22
6,565
3,414
35
6,552
3,414
b
Stem/TA-like (EpiT vs. EpiN)
Stem/TA-like/Im.Goblet (EpiT vs. EpiN)
Stem/TA-like/Prolif. (EpiT vs. EpiN)
20
0
19
0
2
3
30
c
High TFF1
Low TFF1
p = 4.2 x $10^{-2}$ (logrank test)
p = 9.2 x $10^{-2}$ (Cox PH)
n = 219
n = 219
Overall survival
Days
d
No. of patients
cE02_H
cE02_L
cE03_H
cE03_L
e
cE02_H-cE03_H (n = 91)
cE02_L-cE03_H (n =1 26)
cE02_H-cE03_L (n =1 26)
cE02_L-cE03_L (n = 92)
Overall survival
Days
cE02_H-cE03_H vs. rest: HR = 0.62 (0.37 - 1.06), p = 8.0 x $10^{-2}$
cE02_L-cE03_L vs. rest: HR = 1.31 (0.81 - 2.11), p = 2.7 x $10^{-2}$
f
cE02_H-cE03_H (n=122)
cE02_H-cE03_L (n=159)
cE02_L-cE03_H (n=159)
cE02_L-cE03_L (n=122)
Overall survival
Months
cE02_H-cE03_H vs. rest: HR = 0.50 (0.33 - 0.75), p = 8.0 x $10^{-4}$
cE02_L-cE03_L vs. rest: HR = 1.34 (0.97 - 1.85), p = 7.2 x $10^{-2}$
g
cE02_H-cE03_H (n = 120)
cE02_L-cE03_H (n = 158)
cE02_H-cE03_L (n = 157)
cE02_L-cE03_L (n = 122)
Progression-free survival
Months
cE02_H-cE03_H vs. rest: HR = 0.48 (0.31 - 0.73), p = 7.0 x $10^{-4}$
cE02_L-cE03_L vs. rest: HR = 1.47 (1.06 - 2.05), p = 2.0 x $10^{-2}$

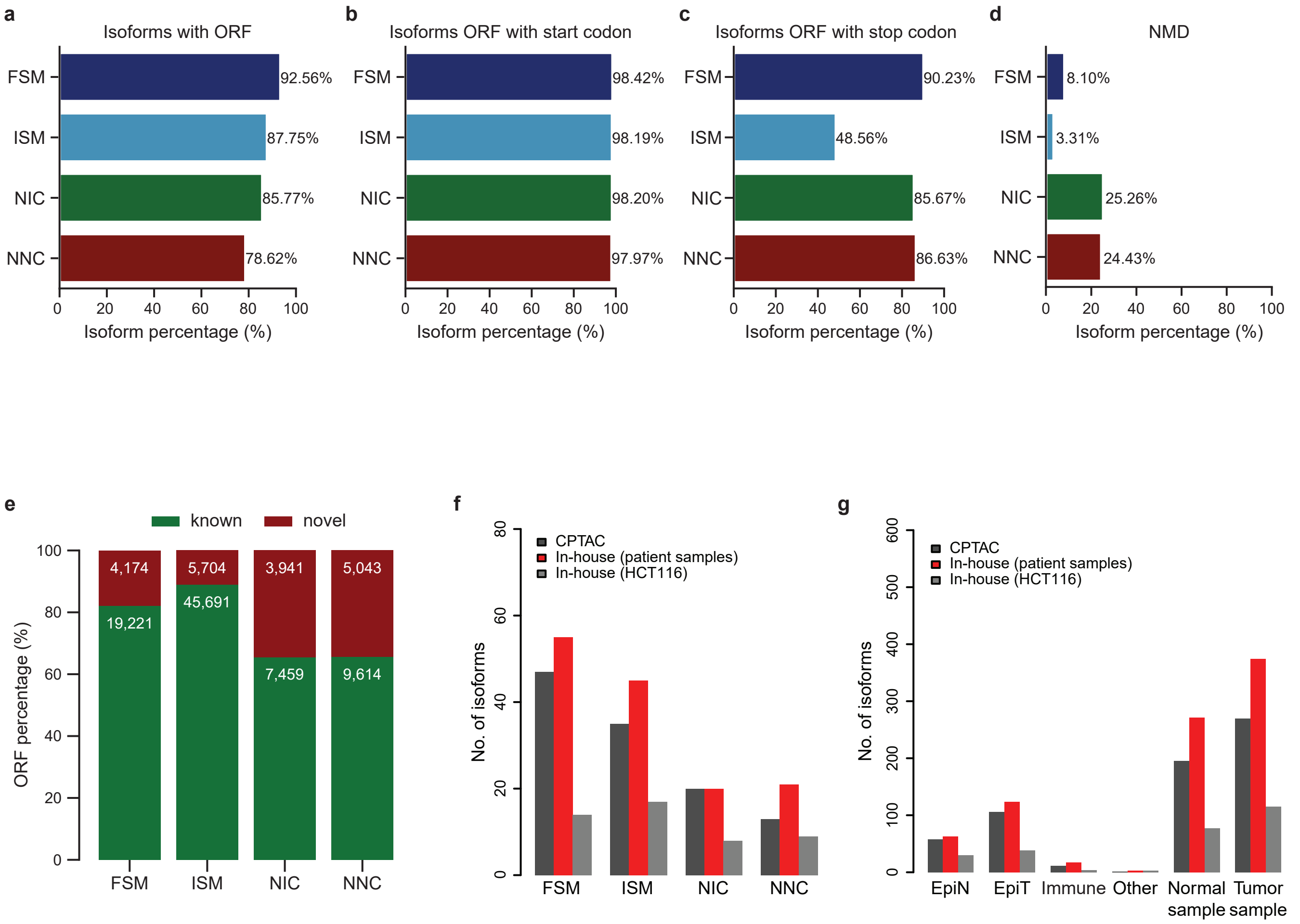
a
Isoforms with ORF
FSM
ISM
NIC
NNC
92.56%
87.75%
85.77%
78.62%
0 20 40 60 80 100
Isoform percentage (%)
b
Isoforms ORF with start codon
98.42%
98.19%
98.20%
97.97%
c
Isoforms ORF with stop codon
90.23%
48.56%
85.67%
86.63%
d
NMD
8.10%
3.31%
25.26%
24.43%
e
known
novel
ORF percentage (%)
4,174
5,704
3,941
5,043
19,221
45,691
7,459
9,614
f
CPTAC
In-house (patient samples)
In-house (HCT116)
No. of isoforms
g
No. of isoforms
EpiN
EpiT
Immune
Other
Normal sample
Tumor sample

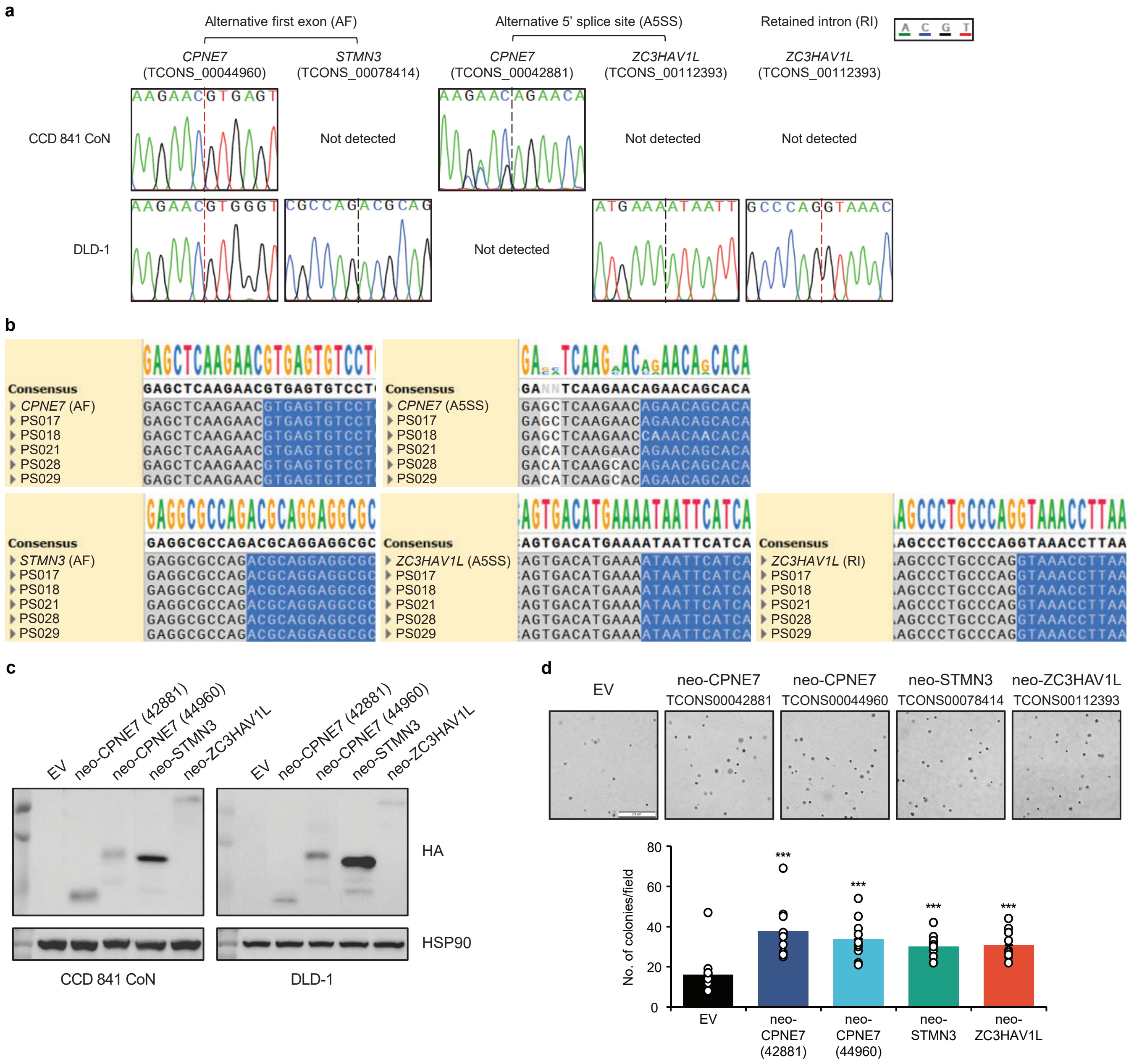


Extended Data Figure 10

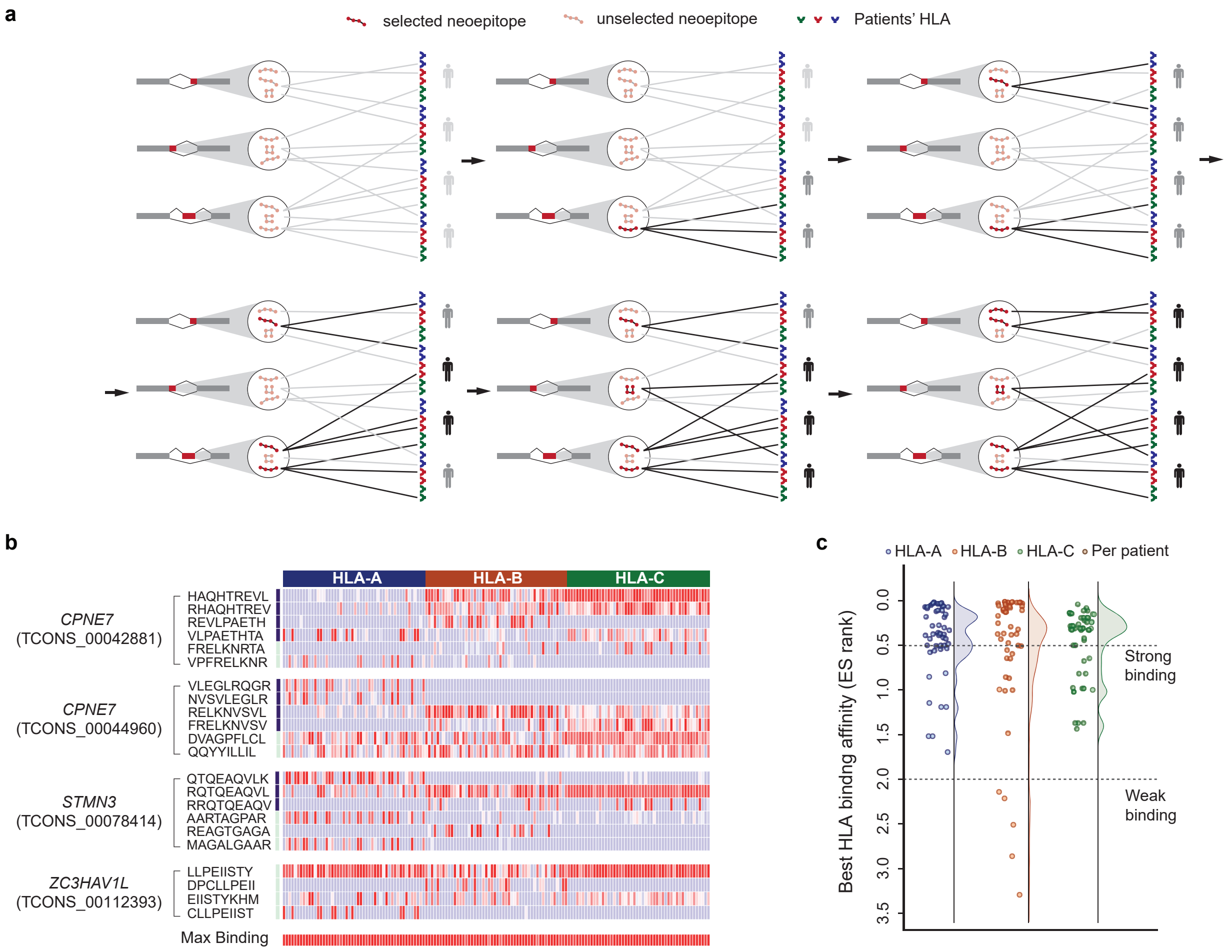
a
selected neoepitope
unselected neoepitope
Patients' HLA
b
HLA-A
HLA-B
HLA-C
CPNE7
(TCONS_00042881)
HAQHTREVL
RHAQHTREV
REVLPAETH
VLPAETHTA
FRELKNRTA
VPFRELKNR
CPNE7
(TCONS_00044960)
VLEGLRQGR
NVSVLEGLR
RELKNVSVL
FRELKNVSV
DVAGPFLCL
QQYYILLIL
STMN3
(TCONS_00078414)
QTQEAQVLK
RQTQEAQVL
RRQTQEAQV
AARTAGPAR
REAGTGAGA
MAGALGAAR
ZC3HAV1L
(TCONS_00112393)
LLPEIISTY
DPCLLPEII
EIISTYKHM
CLLPEIIST
Max Binding
c
HLA-A
HLA-B
HLA-C
Per patient
Best HLA binding affinity (ES rank)
0.0
0.5
1.0
1.5
2.0
2.5
3.0
3.5
Strong binding
Weak binding

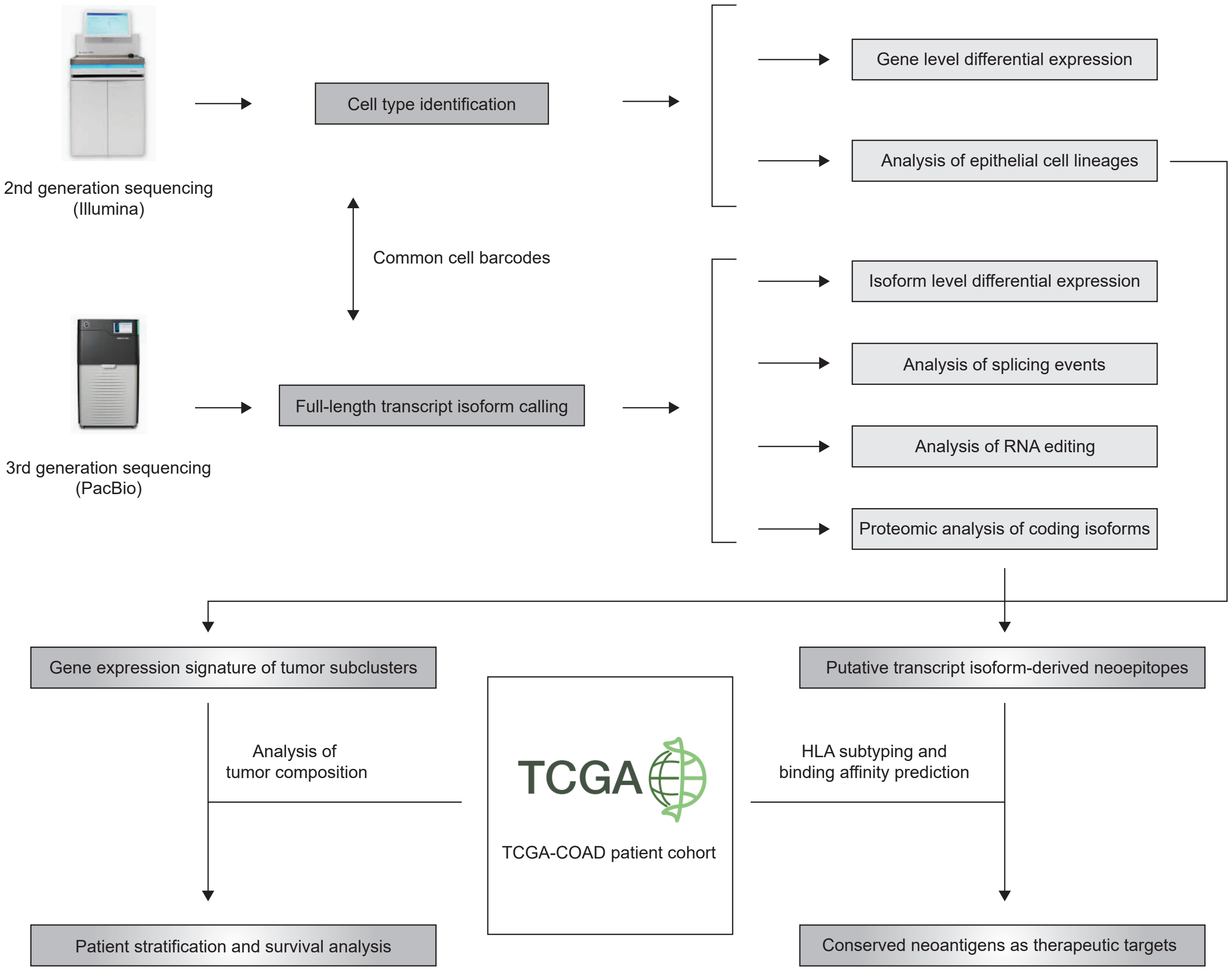
2nd generation sequencing
(Illumina)
Cell type identification
Common cell barcodes
3rd generation sequencing
(PacBio)
Full-length transcript isoform calling
Gene level differential expression
Analysis of epithelial cell lineages
Isoform level differential expression
Analysis of splicing events
Analysis of RNA editing
Proteomic analysis of coding isoforms
Gene expression signature of tumor subclusters
Putative transcript isoform-derived neoepitopes
Analysis of
tumor composition
TCGA
TCGA-COAD patient cohort
HLA subtyping and
binding affinity prediction
Patient stratification and survival analysis
Conserved neoantigens as therapeutic targets
